# Optimized intramuscular immunization with VSV-vectored spike protein triggers a superior protective humoral immune response to SARS-CoV-2

**DOI:** 10.1101/2022.06.14.495413

**Authors:** Adriano Taddeo, Inês Berenguer Veiga, Christelle Devisme, Renate Boss, Philippe Plattet, Sebastian Weigang, Georg Kochs, Volker Thiel, Charaf Benarafa, Gert Zimmer

## Abstract

Immunization with vesicular stomatitis virus (VSV)-vectored COVID-19 vaccine candidates expressing the SARS-CoV-2 spike protein in place of the VSV glycoprotein relies implicitly on expression of the ACE2 receptor at the muscular injection site. Here, we report that such a viral vector vaccine did not induce protective immunity following intramuscular immunization of K18-hACE2 transgenic mice. However, when the viral vector was *trans*-complemented with the VSV glycoprotein, intramuscular immunization resulted in high titers of spike-specific neutralizing antibodies. The vaccinated animals were fully protected following infection with a lethal dose of SARS-CoV-2-S^D614G^ via the nasal route, and partially protected if challenged with the SARS-CoV-2^Delta^ variant. While dissemination of the challenge virus to the brain was completely inhibited, replication in the lung with consequent lung pathology was not entirely controlled. Thus, intramuscular immunization was clearly enhanced by *trans*-complementation of the VSV-vectored vaccines by the VSV glycoprotein and led to protection from COVID-19, although not achieving sterilizing immunity.

## Introduction

Vesicular stomatitis virus (VSV) is a non-segmented negative-strand RNA virus, which belongs to the family of *Rhabdoviridae*. Genetically engineered VSV expressing foreign antigens have been widely used as vector vaccines for protection against several human and veterinary pathogens.^1, 2^ The VSV-based Ebola vaccine, which has been approved for human use, represents a replication-competent vector in which the single type VSV envelope glycoprotein G was replaced by the Ebola virus glycoprotein.^3^ A similar approach was used to generate VSV-vectored vaccines for protection from coronavirus disease 2019 (COVID-19). The VSV envelope glycoprotein G was replaced with the SARS-CoV-2 spike antigen resulting in a replication-competent chimeric virus.^4, 5^ The vaccine candidates showed promising results in preclinical mouse and hamster animal models. A single intramuscular immunization was already sufficient to induce high titers of neutralizing antibodies and to protect the animals from SARS-CoV-2 challenge infection.^4, 5^ Based on these encouraging results, phase 1 clinical studies were performed with these vaccine candidates in Israel (ClinicalTrials.gov Identifier: NCT04608305) and the US (ClinicalTrials.gov Identifier: NCT04569786), but were not continued with clinical phase 2/3 trials. Unfortunately, the results of these clinical studies have not been disclosed, and the reasons why further evaluation of the vaccine candidates was abandoned have not been disclosed.

SARS-CoV-2 primarily replicates in epithelial cells of the respiratory tract.^6^ The infection is initiated by binding of the viral spike (S) protein to the cellular receptor angiotensin converting enzyme 2 (ACE2).^7^ Following primary replication of SARS-CoV-2 in the respiratory epithelium, the virus can disseminate to extrapulmonary tissues such as the vascular endothelium, heart, kidney, intestine and the neuronal system.^8–11^ In line with this broad tropism of SARS-CoV-2, the ACE2 receptor has been detected in various cell types, however, the expression of ACE2 by skeletal muscle cells has not been reported.^12–14^ We suppose that the intramuscular vaccination of humans with the VSV-vectored vaccine candidates in recent clinical phase 1 studies might have resulted in only low infection rates and therefore inefficient stimulation of the immune system.

Here we generated a propagation-competent viral vector, VSV*ΔG-S_Δ21,_ by substituting the VSV glycoprotein G with the SARS-CoV-2 spike protein. We demonstrate that *trans*-complementation of VSV*ΔG-S_Δ21_ with the VSV G glycoprotein allowed infection of cells lacking the ACE2 receptor, and that ACE2-independent immunization was superior over ACE2-dependent immunization in inducing neutralizing antibodies and protective immunity in mice. Importantly, intramuscular immunization did not prevent SARS-CoV-2 replication in the respiratory tract and consequent lung pathology, particularly when using a challenge virus that did not completely match the vaccine antigen.

## Results

### Generation of VSV-vectored vaccine candidates

The VSV vector vaccine VSV*ΔG-S_Δ21_ was generated by replacing the glycoprotein (G) gene in the VSV genome with cDNA encoding a modified SARS-CoV-2 spike (S) protein truncated at the C-terminus by 21 amino acids (Δ21) to improve incorporation into the VSV envelope.^15^ The S1/S2 proteolytic cleavage site of the S protein was modified by introducing the mutation R685G, which reduces S-mediated syncytia formation and enhances virus yield *in vitro*.^4, 5^ An additional transcription unit encoding the green fluorescent protein (GFP) was inserted into the vector genome downstream of the S gene (**Fig. 1a**). A second vaccine candidate, VSV*M_q_ΔG-S_Δ21_, was generated by replacing the matrix (M) gene in the VSV*ΔG-S_Δ21_ vector genome with the modified M_q_ gene, which encodes an M protein devoid of host shut-off activity.^16^ Finally, a propagation-defective vector vaccine, VSV*ΔG-G_RBD_, was designed by inserting the synthetic G_RBD_ gene into the VSV vector genome. G_RBD_ encodes a composite membrane protein made up of the N-terminal VSV G signal peptide, the receptor-binding domain (RBD) of the SARS-CoV-2 S protein, and the C-terminal part of the VSV G protein comprising the VSV G stem, transmembrane, and cytosolic domains.

**Fig. 1.**
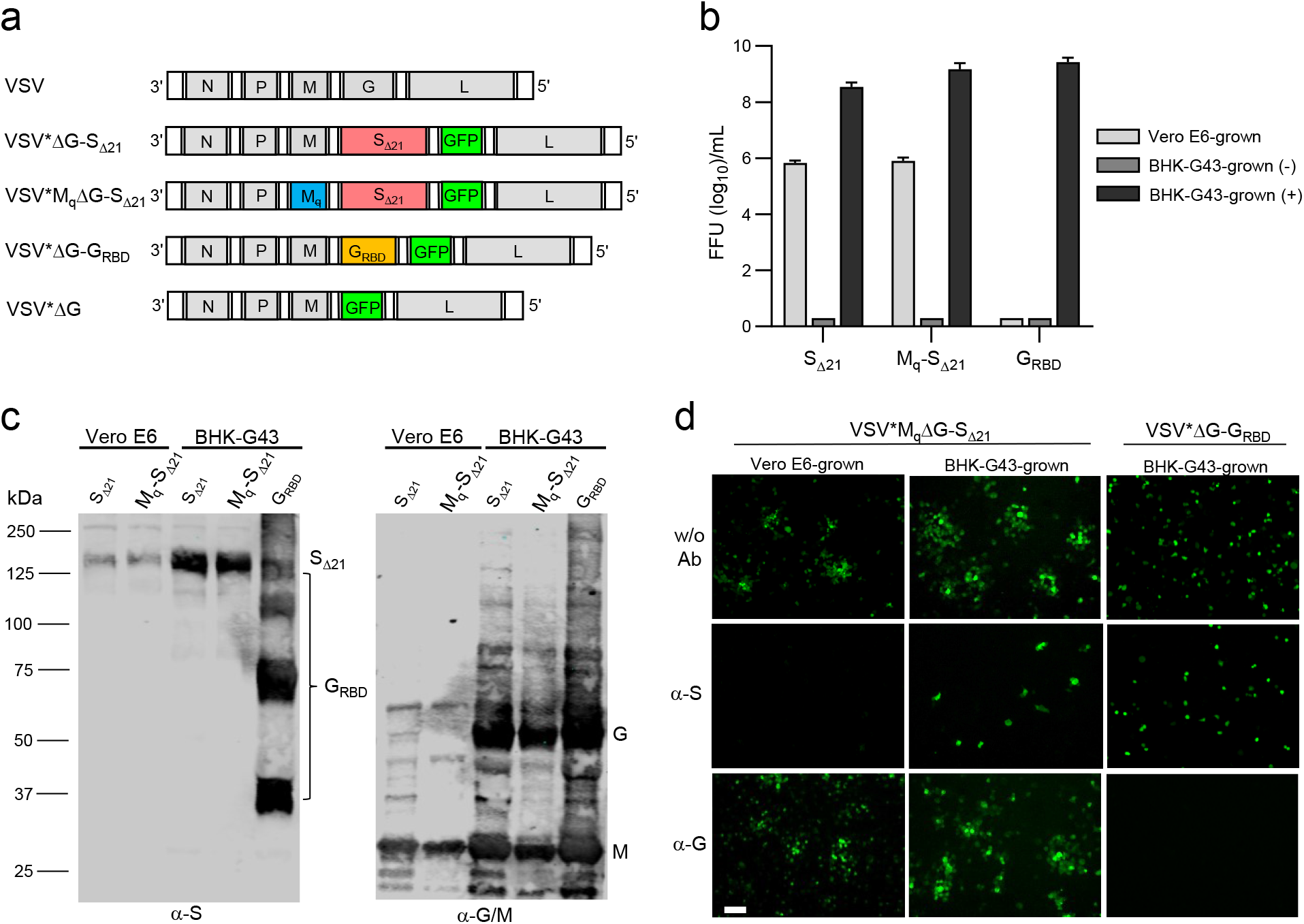
Generation and characterization of VSV-vectored COVID-19 vaccine candidates. (**a**) Genome maps of recombinant VSV vectors. The original VSV contains five transcription units encoding the N, P, M, G, and L genes. The VSV vector was modified by replacing the G gene with a modified SARS-CoV-2 spike gene and by inserting an additional transcription unit encoding GFP. M_q_ denotes a quadruple mutant M gene encoding an M protein that lacks host shut-off activity. G_RBD_ denotes a synthetic membrane protein presenting the receptor-binding domain (RBD) of the SARS-CoV-2 spike protein. (**b**) Virus yield on Vero E6 and BHK-G43 cells. Vero E6 cells, BHK-G43 cells expressing the VSV glycoprotein (+), and BHK-G43 cells lacking VSV glycoprotein expression (-) were infected with either VSV*M_q_ΔG-S_Δ21_ (S_Δ21_), VSV*M_q_ΔG-S_Δ21_ (M_q_-S_Δ21_), or VSV*ΔG-G_RBD_ (G_RBD_) using 0.1 ffu/cell. At the 24 hours pi, infectious virus released into the cell culture was titrated on Vero E6 cells. Mean values and standard deviations of 3 infection experiments are shown. (**c**) Western blot analysis of recombinant VSV vector particles. VSV*ΔG-S_Δ21_ (S_Δ21_) and VSV*M_q_ΔG-S_Δ21_ (M_q_-S_Δ21_) were propagated on both Vero E6 and BHK-G43 cells, while VSV*ΔG-G_RBD_ (G_RBD_) were propagated only on BHK-G43 helper cells. At 24 hours pi, the virus particles were concentrated from the cell culture supernatant by ultracentrifugation and dissolved in SDS sample buffer. The viral proteins were separated by SDS-PAGE under non-reducing conditions and blotted onto nitrocellulose membrane. Antigens were detected with a COVID-19 convalescent serum (α-S) and a rabbit polyclonal immune serum directed to the VSV G and M proteins (α-G/M). The blots were derived from the same experiment and were not further processed. (**d**) Inhibition of virus entry using neutralizing antibodies directed to either the VSV G protein or the SARS-CoV-2 spike protein. Vero E6 cells were inoculated in the absence or presence of the indicated neutralizing antibodies with VSV*M_q_ΔG-S_Δ21_ produced on either Vero E6 or BHK-G43 cells and with VSV*ΔG-G_RBD_ grown on BHK-G43 cells. Infected cells were detected 24 hours pi taking advantage of the GFP reporter. Bar = 100 µm.

BHK-G43 cells are transgenic BHK-21 cells, which express the VSV G protein in a regulated manner.^17^ All three VSV-vectored vaccines, VSV*ΔG-S_Δ21_, VSV*M_q_ΔG-S_Δ21_, and VSV*ΔG-G_RBD_ replicated on these cells to high titers if expression of the VSV glycoprotein was induced (+), but none of them propagated on them in the absence of VSV glycoprotein expression (-) (**Fig. 1b**). VSV*ΔG-S_Δ21_ and VSV*M_q_ΔG-S_Δ21_ also replicated on Vero E6 cells, however, titers were approximately 1000-fold lower than those on Vero E6 cells.

In A549 cells stably expressing both human ACE2 and human TMPRSS2, VSV*ΔG-S_Δ21_ replicated slowly but reached titers of about 10^6^ ffu/ml at 72 hours post infection (pi) (**Supplementary Fig. 1a**). In contrast, VSV*M_q_ΔG-S_Δ21_ reached only 10^2^ ffu/ml at 72 hours pi, suggesting that the mutant M_q_ protein limited virus replication in this interferon-competent cell line. Accordingly, IFN bioactivity was detected in the supernatant of A549 cells infected with VSV*M_q_ΔG-S_Δ21_, but not in the supernatant of VSV*ΔG-S_Δ21_-infected cells (**Supplementary Fig. 1b**). These results indicate that the VSV*M_q_ΔG-S_Δ21_ vector is propagation-restricted in IFN-competent cells.

The propagation of the vaccine candidates on either Vero E6 or BHK-G43 helper cells was expected to affect the composition of the viral envelope and consequently the susceptibility of the viral vector to neutralizing antibodies. The VSV M protein was present in all VSV vaccine species grown in BHK-G43 or Vero E6 cells, while the VSV G protein was expressed only in the BHK-G43-grown viruses (**Fig. 1c,** α-G/M blot). The S_Δ21_ protein was detected with an apparent molecular mass of 150 kDa under non-reducing conditions in both VSV*ΔG-S_Δ21_ and VSV*M_q_ΔG-S_Δ21_ grown on both host cells (**Fig. 1c,** α-S blot). Under reducing conditions, the S_Δ21_ protein signal was reduced, probably due to disruption of conformation-dependent epitopes (**Supplementary Fig. 1c**, α-S blot). The synthetic G_RBD_ protein appeared as several protein bands at 36, 72, 110, and 140 kDa (**Fig. 1c,** α-S blot). The 36 kDa band corresponds to the predicted molecular mass of the G_RBD_ protein, while the higher molecular weight bands may represent oligomeric forms of G_RBD_. When SDS-PAGE was run under reducing conditions, the 36 kDa band was primarily detected suggesting that the putative oligomeric forms of the protein are linked by disulfide bonds (**Supplementary Fig. 1c,** α-S blot). Functionally, infection of Vero E6 cells with VSV*M_q_ΔG-S_Δ21_ grown on Vero E6 cells was completely inhibited by COVID-19 convalescent serum (α-S), whereas a neutralizing antibody directed to the VSV G protein (α-G) had no effect (**Fig. 1d, left panels**). Infection of Vero E6 cells with VSV*M_q_ΔG-S_Δ21_ grown on BHK-G43 (expressing both VSV G and SARS-CoV-2 S) was neither blocked by α-G nor α-S, but α-S prevented further spreading of the virus in the cell monolayer (**Fig. 1d, central panels**). Finally, infection of Vero E6 cells with VSV*ΔG-G_RBD_ grown on BHK-G43 led to the formation of single-cell foci, which were completely blocked by α-G serum, while α-S had expectedly no effect (**Fig.1d, right panels**).

### VSV G *trans*-complementation of VSV*ΔG-S_Δ21_ increases the immune response to spike antigen

To compare ACE2-dependent with ACE2–independent immunization, VSV*ΔG-S_Δ21_ was propagated on Vero E6 and BHK-G43 cells, respectively, and then injected intramuscularly in groups of 5 K18-hACE2 mice using 10^4^ or 10^5^ ffu per animal. Serum was collected 3 weeks after the first (prime) and the second (boost) injections, which were performed at a 4-week interval. Immunization with BHK-G43-grown VSV*ΔG-S_Δ21_ induced spike-specific antibodies at significantly higher levels than the Vero E6 cell-grown vector using an anti-S ELISA (**Fig. 2a**). ELISA titers were neither significantly increased when using a higher dose nor after the boost. In contrast, the BHK-G43 grown vector induced neutralizing antibodies in a dose-dependent manner, and neutralization titers were significantly increased after the second immunization (**Fig. 2b**). Notably, Vero E6-grown VSV*ΔG-S_Δ21_ did not induce any virus-neutralizing activity. C57BL/6J mice that do not express hACE2 receptor showed a similar serological profile as K18-hACE2 mice after immunization with the same VSV*ΔG-S_Δ21_ vector preparations (**Supplementary Fig. 2a, b**). Immunization of K18-hACE2 mice with 10^6^ ffu of the BHK-G43-grown control vaccine VSV*ΔG did not trigger any spike-specific antibodies as measured by ELISA (**Sup. Fig. 3a**) and virus neutralization test (**Fig. 3a**).

**Fig. 2.**
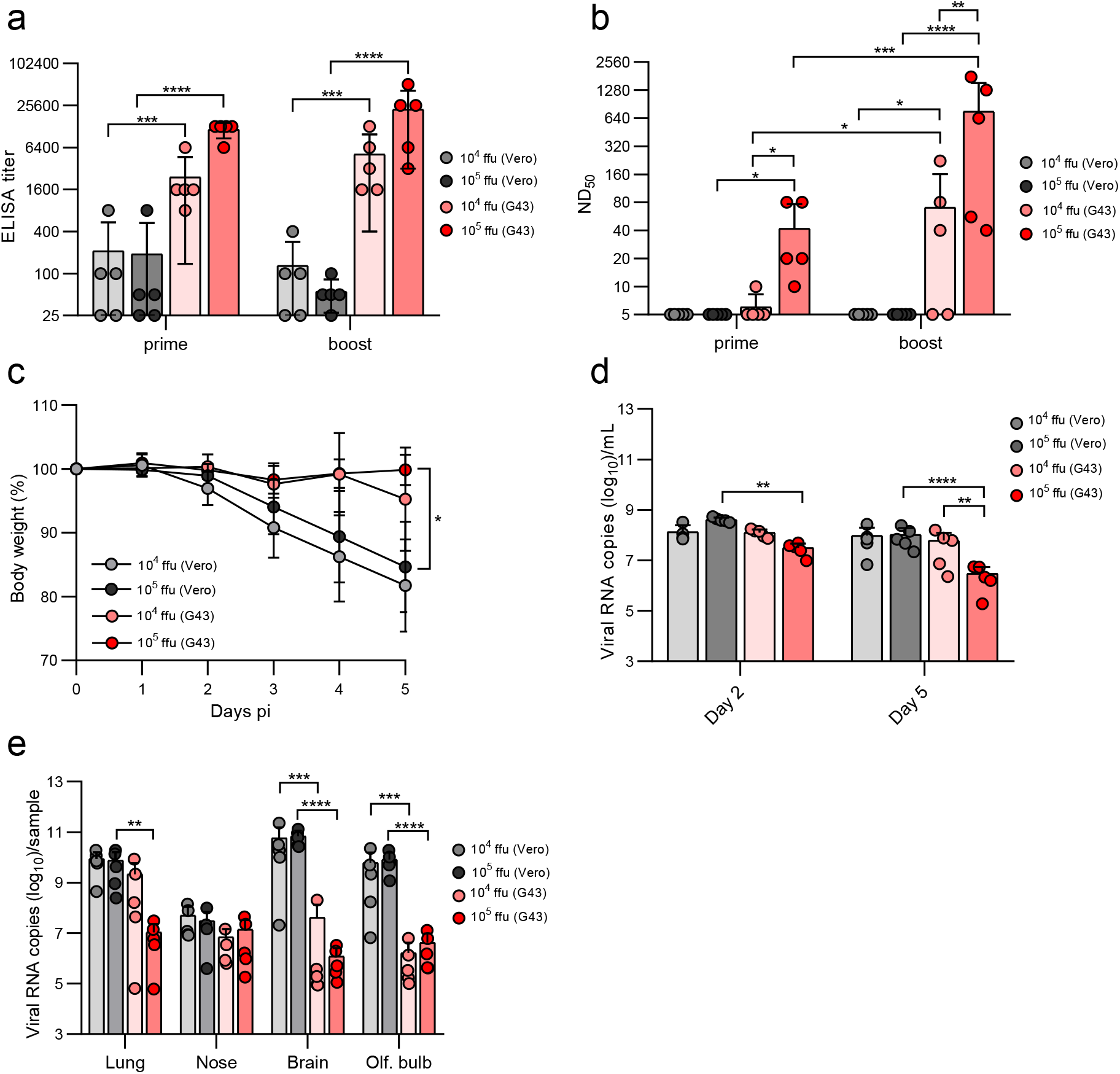
Comparison of ACE2-dependent and -independent immunization. K18-hACE2 mice were immunized (i.m.) with either 10^4^ or 10^5^ focus-forming units (ffu) of the VSV*ΔG-S_Δ21_ vector vaccine which was produced on either Vero E6 or BHK-G43 cells. (**a**) Detection of spike-specific serum antibodies by ELISA 3 weeks after the first (prime) and 3 weeks after the second immunization (boost). (**b**) Determination of the virus neutralization dose 50% (ND_50_) in serum of immunized mice. (**c**) Determination of body weight of immunized mice following nasal infection with 10^5^ pfu of SARS-CoV-2-S^D614G^. (**d**) Determination of virus load by RT-qPCR in oropharyngeal swab samples collected at days 2 and 5 pi and (**e**) in the indicated organs prepared from the euthanized animals at day 5 pi. Mean values and standard deviations for n = 5 mice per group are shown. Statistical analysis was performed using the two-way ANOVA with either Tukey’s or Sidak’s multiple comparison test (*p < 0.05; **p < 0.005; ***p < 0.0005; ****p < 0.0001).

**Fig. 3.**
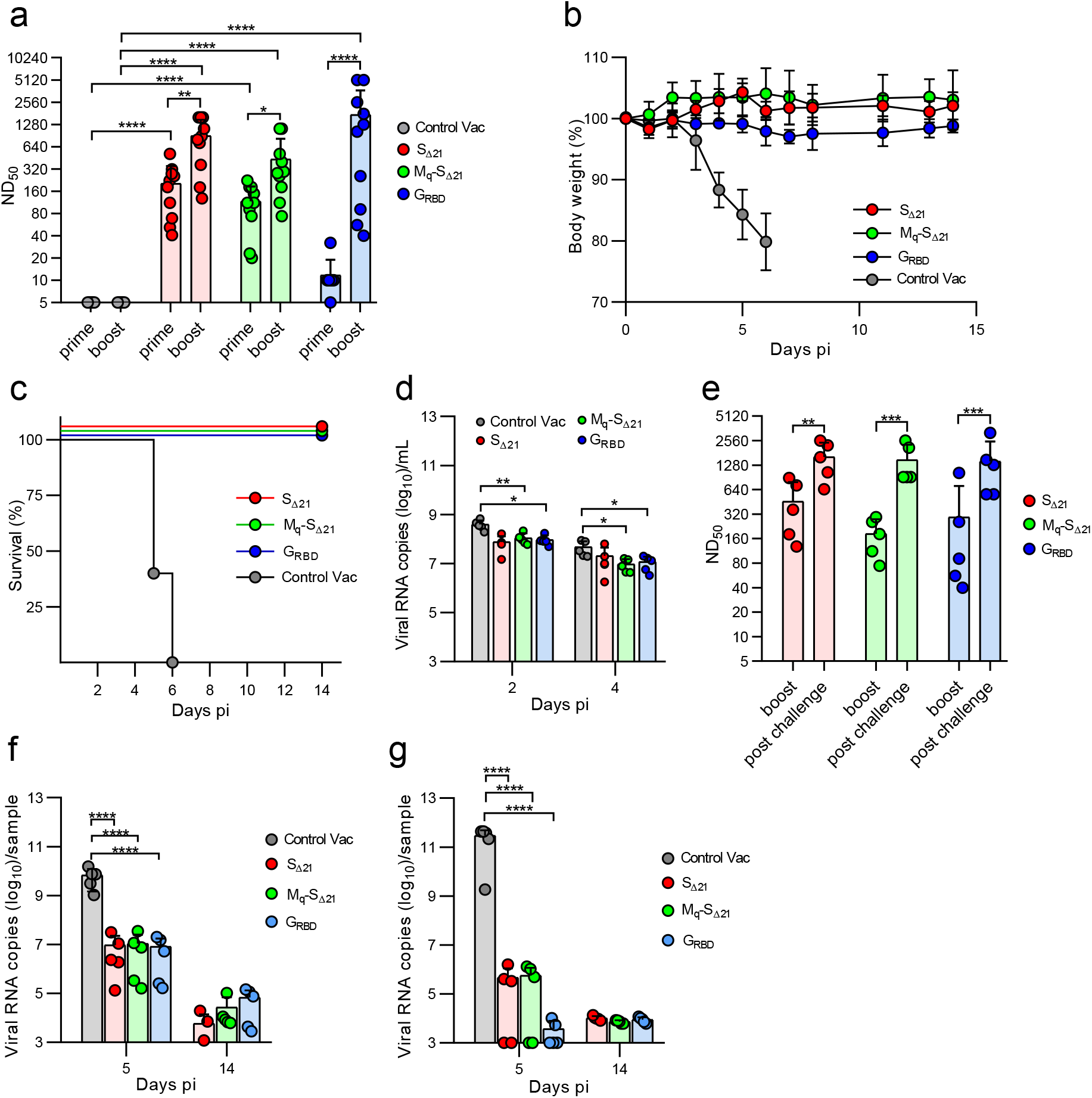
Analysis of protection of immunized K18-hACE2 mice from SARS-CoV-2 challenge infection. K18-hACE2 mice were immunized twice (i.m.) with one of the following vaccines: VSV*ΔG-S_Δ21_ (n = 10), VSV*M_q_ΔG-S_Δ21_ (n = 10), VSV*ΔG-G_RBD_ (n = 10), and the VSV*ΔG control vaccine (n = 5). Five animals of each vaccine group were euthanized at day 5 pi, while the others were maintained until day 14 pi. (**a**) Determination of neutralizing serum antibody titers (ND_50_) 21 days after the primary immunization (prime) and 18 days after the secondary immunization (boost). Weight loss (**b**) and survival (**c**) of K18-hACE2 mice (d14 group) following nasal infection with 10^5^ pfu of SARS-CoV-2-S^D614G^. (**d**) Quantification (RT-qPCR) of SARS-CoV-2 RNA copies in oropharyngeal swab samples collected at 2 and 4 days pi. (**e**) Determination of neutralizing serum antibody titers (ND_50_) 18 days after the boost and 14 days post challenge infection. (**f, g**) RT-qPCR determination of viral genome copy numbers in lung (**f**) or brain (**g**) homogenates prepared from animals that were euthanized at either day 5 or day 14 pi. Mean values and standard deviations (SD) are shown. Statistically significant differences as computed by either the two-way ANOVA test (**a, d, e**) or the one-way ANOVA test (**f, g**) are indicated (*p < 0.05; **p < 0.005; ***p< 0.0005; ****p < 0.0001).

Four weeks after the boost, the four groups of immunized K18-hACE2 mice were challenged with a SARS-CoV-2 strain encoding the S(D614G) mutation (SARS-CoV-2-S^D614G^). While mice immunized with Vero E6-grown VSV*ΔG-S_Δ21_ virus lost body weight (**Fig. 2c**) to a similar degree than mice immunized with the control vaccine (**Fig. 3b**), mice immunized with BHK-G43-grown virus were protected against weight loss until the termination of the experiment at day 5 pi (**Fig. 2c**). No effect of vector vaccine dose on weight loss was observed. Virus load in oropharyngeal swabs were significantly reduced in mice immunized with 10^5^ ffu BHK-G43-grown vector compared to mice that received 10^4^ ffu (day 5 pi) and to those receiving the Vero E6-grown vector (days 2 and 5 pi) (**Fig. 2d**). In the lungs of mice vaccinated with either 10^4^ or 10^5^ ffu of Vero E6-grown VSV*ΔG-S_Δ21,_ the SARS-CoV-2 virus load at day 5 pi was about 10^10^ viral RNA genomic copies/sample (**Fig. 2e**), which corresponded to the virus load detected in the lungs of control-vaccinated mice (**Fig. 3f**). In contrast, vaccination with 10^5^ ffu, but not with 10^4^ ffu, of BHK-G43-grown vector caused a significant reduction of SARS-CoV-2-S^D614G^ load in the lung measured by qPCR **(Fig. 2e)**, while both doses of this vaccine resulted in the absence of detectable infectious virus particles by plaque assay (**Supplementary Fig. 2c;** compare with **Supplementary Fig. 3d** showing the number of virus plaques found in the lungs of control-vaccinated mice). The virus loads were reduced in the brain and olfactory bulb, but not in the nose, of mice immunized with BHK-G43-grown vaccine doses compared to the corresponding Vero E6-grown vaccines **(Fig. 2e,** see also **Fig. 3g** showing the virus loads in the brain of control-vaccinated mice). No infectious virus was detected in the brain of all but one mouse immunized with BHK-G43-grown vaccine, while nine out of 10 mice immunized with Vero E6-grown vaccines had high PFUs in the brain **(Supplementary Fig. 2d;** see also **Supplementary Fig. 3e** for infectious virus in the brains of control-vaccinated mice). Together, the findings show the superiority of *trans*-complemented vector in the induction of a protective immunity against severe systemic disease in K18-hACE2 mice.

### Evaluation of VSV-vectored vaccines in the K18-hACE2 mouse model

K18-hACE2 mice were immunized intramuscularly with 2x 10^6^ ffu of either VSV*ΔG-S_Δ21_, VSV*M_q_ΔG-S_Δ21_, or VSV*ΔG-G_RBD_ using a prime-boost protocol with a 4-week interval. A fourth group received the VSV*ΔG vector as control vaccine (Control Vac), which did not encode any SARS-CoV-2 antigen (**Fig. 1a**). All vaccine candidates were produced on BHK-G43 cells and therefore *trans*-complemented with the VSV G protein. Serum was collected 25 days after the prime and 18 days after the boost vaccination and analyzed for the presence of virus-neutralizing antibodies (**Fig. 3a**). A single immunization with VSV*ΔG-S_Δ21_ and VSV*M_q_ΔG-S_Δ21_ resulted in the induction of high titers of neutralizing antibodies (mean ND_50_ value of 204 and 116, respectively), whereas the propagation-defective VSV*ΔG-G_RBD_ vector induced neutralizing antibodies at very low levels (mean ND_50_ value of 12), which did not significantly differ from ND_50_ values in control-vaccinated mice. However, the titers increased 144-fold in this vaccine group following the second immunization (mean ND_50_ value of 1732). The VSV*ΔG-S_Δ21_ and VSV*M_q_ΔG-S_Δ21_ vaccine groups also showed significantly increased neutralizing antibody levels after the boost, although the increase was not that dramatically (4.4-fold and 3.8-fold, respectively). The immune sera of 5 animals of each vaccine group were also analyzed for the presence of spike-specific antibodies by ELISA (**Supplementary Fig. 3a)**. In addition, the frequency of circulating CD8+ T cells binding H-2K(b) MHC tetramers loaded with the S 539-546 peptide was analyzed by flow cytometry 4 weeks after the boost. Compared to naïve animals, an increased frequency of S-specific CD8+ T cells were found in the VSV*ΔG-S_Δ21_ and VSV*ΔG-G_RBD_ vaccine groups (**Supplementary Fig. 3b, c**), indicating that VSV-vectored vaccines have the potential to trigger CD8^+^ T cell immune responses.

Immunized K18-hACE2 mice were intranasally infected with 10^5^ plaque-forming units (pfu) of SARS-CoV-2-S^D614G^ 3 weeks after the boost and 5 mice of each vaccine group were assigned to be euthanized at day 5 and day 14 pi. The animals of the control group rapidly lost weight (**Fig. 3b**) and reached exclusion criteria at days 5 and 6 pi (**Fig. 3c**). In contrast, all animals vaccinated with either of the three vaccine candidates survived the infection with SARS-CoV-2 without showing any body weight loss (**Fig. 3b, c**). RT-qPCR analysis of oropharyngeal swab samples collected at days 2 and 4 pi revealed that virus load was significantly reduced in the VSV*M_q_ΔG-S_Δ21_, and VSV*ΔG-G_RBD_ vaccine groups compared to the Control Vac group (**Fig. 3d**). The reduction of virus loads in swab samples of the VSV*ΔG-S_Δ21_ group did not reach significance due to high variability. High neutralizing antibody titers with mean ND_50_ values of 1620, 1479, and 1418, were detected at day 14 post challenge in sera of the VSV*ΔG-S_Δ21_, VSV*M_q_ΔG-S_Δ21_, and VSV*ΔG-G_RBD_ vaccine groups, respectively (**Fig. 3e**), indicating that the challenge infection further boosted the spike-specific immune response.

### VSV-vectored vaccines reduce SARS-CoV-2 replication in the lung and block virus dissemination to the brain

Analysis of SARS-CoV-2 genomic RNA levels in lung homogenates at day 5 pi revealed a reduction of virus load by 3 log_10_ for all three vector vaccines compared to the control vaccine (**Fig. 3f)**. SARS-CoV-2 load in the lungs of infected mice was further reduced at day 14 pi to levels close to the detection limit of 10^3^ viral RNA copies/ml. At day 5 pi, significantly higher levels of SARS-CoV-2 genomic RNA were detected in the brain of the control vaccine group compared to the vaccinated groups, which had 6 log_10_ less virus **(Fig. 3g**). The virus load in brain of the vaccinated animals was further reduced to the detection limit at day 14 pi. Moreover, infectious virus was not detected in lung and brain homogenates of the mice of the 3 vaccinated groups, neither at day 5 nor at day 14 pi; high virus titers were found in the control vaccine group on day 5 pi (**Supplementary Fig. 3d, e**).

Histopathological analysis of the lungs from animals of the control vaccine group, which reached exclusion criteria 5 to 6 days pi (**Fig. 3c**) revealed a moderate to severe lymphohistiocytic interstitial pneumonia, characterized by extensive interstitial, peribronchiolar and perivascular inflammation, which extended into the alveoli with variable degrees of alveolar and perivascular edema (**Fig. 4a, HE Lung, Day 5 pi**). The immunohistochemical (IHC) analysis of the lung of these mice with an antibody specific for the SARS-CoV-2 nucleoprotein revealed a very strong and widespread viral antigen detection, while lung tissue sections of non-infected mice remained negative (**Fig. 4a, IHC Lung**). All animals vaccinated with either VSV*ΔG-S_Δ21_, VSV*M_q_ΔG-S_Δ21_ or VSV*ΔG-G_RBD_ showed variable degrees of inflammation as well as rare vascular changes at day 5 pi, which partially resolved at day 14 pi (**Fig. 4a, HE Lung, Day 14 pi**). IHC analysis of these animals revealed solely a few infectious foci in lung sections corresponding to day 5 pi, which were absent from lung tissue sections corresponding to day 14 pi. Previously, SARS-CoV-2 infection of the brain has been shown to cause the death of K18-hACE2 mice.^18^ Accordingly, IHC analysis of brain sections from day 5 pi revealed widespread SARS-CoV-2 nucleoprotein detection in the control vaccine group (**Fig. 4a, IHC Brain)** in association with very mild lesions (**Fig. 4a, HE Brain**), while no evidence for infection was observed in brain tissue sections from animals of either vaccine groups. Finally, semi-quantitative assessment of lung pathology revealed that mice of the VSV*M_q_ΔG-S_Δ21_ vaccine group had significantly less histopathological changes at day 5 pi compared to the control vaccine group (**Fig. 4b**). However, animals of the vaccine groups still showed significantly enhanced lung pathology scores at day 14 pi compared to lungs from non-infected mice (**Fig. 4c**). Together, these results indicate that intramuscular immunization with VSV-vectored vaccines triggered an immune response that reduced virus replication in the lung and effectively prevented dissemination of SARS-CoV-2 to the brain.

**Fig. 4.**
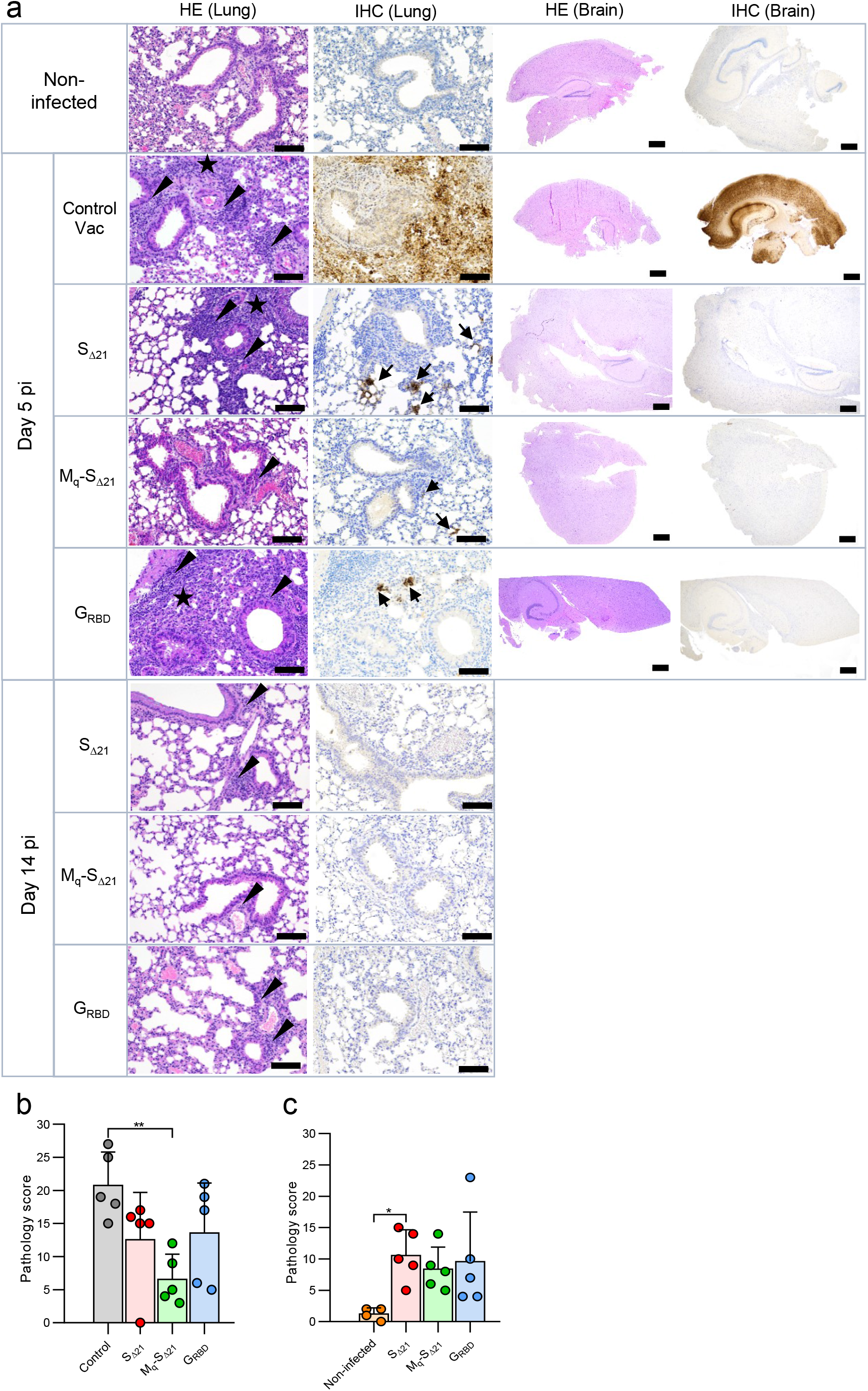
Analysis of lung pathology in SARS-CoV-2 infected K18-hACE2 mice. K18-hACE2 mice (group size n = 10) were immunized with the indicated vaccines and challenged with 10^5^ pfu of SARS-CoV-2-S^D614G^ via the nasal route. Five animal groups of each vaccine group were euthanized at day 5 pi while the remaining animals of each group were sacrificed at day 14 pi. (**a**) Lung and brain tissue sections were analyzed by immunohistochemistry (IHC) using a rabbit polyclonal antibody directed to the SARS-CoV nucleoprotein antigen. Lung and brain sections prepared from mock-infected non-vaccinated animals served as control. Arrows indicate lung and brain areas where the viral nucleoprotein antigen was detected. In parallel, lung and brain tissue sections were stained by hematoxylin-eosin (HE). Arrow heads indicate perivascular/peribronchiolar infiltration. Stars mark tissue consolidation. Large size and small size bars indicate 100 and 500 µm, respectively. (**b, c**) Histopathological scoring of lung tissue sections obtained on days 5 (**b**) and 14 (**c**) pi. Mean values and SD are indicated. Statistically significant differences as computed by the one-way ANOVA test are shown (*p < 0.05; **p < 0.01).

### Maternal antibodies protect offspring from lethal infection with SARS-CoV-2-S^D614G^

We then investigated whether maternal antibodies would protect offspring of immune K18-hACE2 mice from challenge with SARS-CoV-2. K18-hACE2 females vaccinated twice with VSV*M_q_ΔG-S_Δ21_ and challenged with SARS-CoV-2-S^D614G^ were co-housed for 5 days with K18-hACE2 males starting at 1 day pi. Five weeks after birth, virus-neutralizing antibodies were detected (mean ND_50_ of 38) from the pups from such matings, while no neutralizing antibodies were found in from naïve age-matched mice (**Fig. 5a**). One week later, the 6-week-old offspring were infected via the nasal route with 3×10^4^ pfu of SARS-CoV-2-S^D614G^ and euthanized on day 5 pi to determine virus and antibody titers. While offspring of naïve mice lost weight, 4 out of 5 offspring from vaccinated/infected mice gained weight (**Fig. 5b**). One offspring of the immune mothers lost weight and had correspondingly no more detectable neutralizing activity in serum post challenge (**Fig. 5a**) and high virus titers in the brain on day 5 pi (**Fig. 5c**). At that time point, the four protected offspring had virus-neutralizing antibody at levels that were in the same range as measured two weeks before (**Fig. 5a**), and virus titers were found in lungs and brain (**Fig. 5c**). Offspring of naïve mice had virus titers in lungs and brain. These results indicate that passive transfer of maternal antibodies protect mice from lethal SARS-CoV-2 infection.

**Fig. 5.**
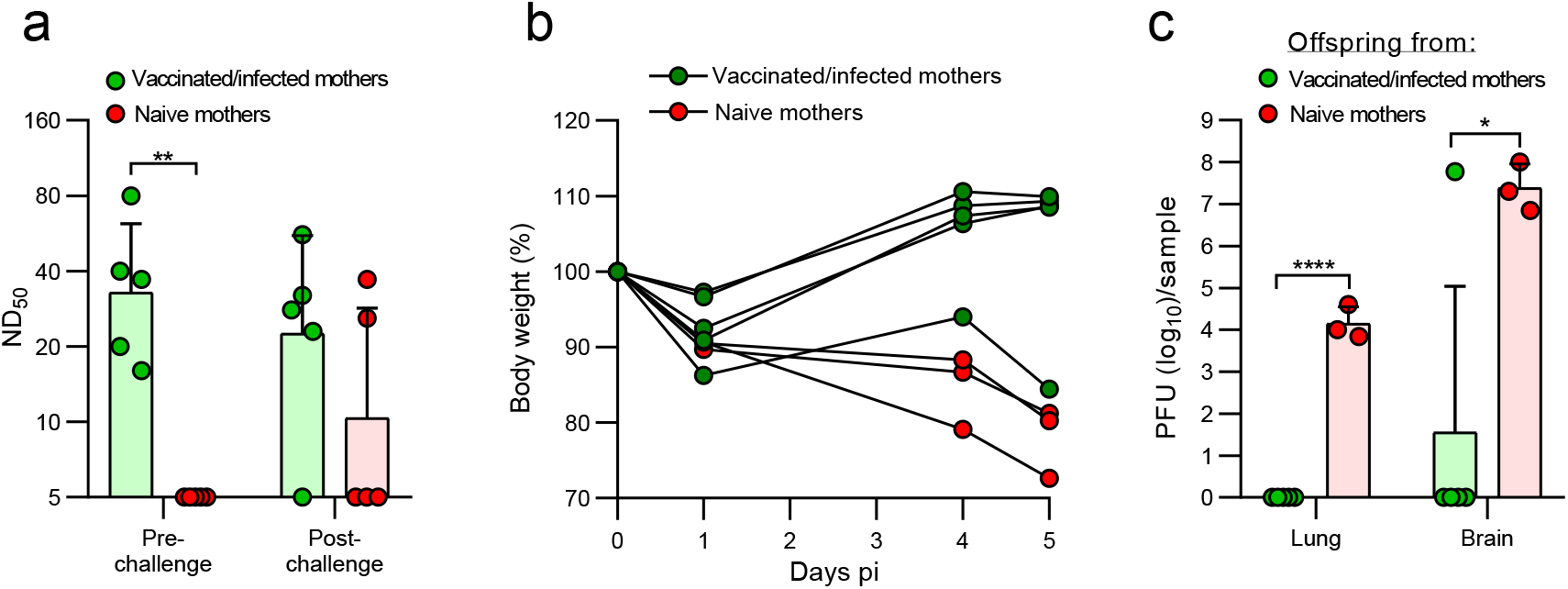
Maternal antibodies protect offspring from lethal SARS-CoV-2 infection. Six-week-old offspring of naïve (red) or of immunized and subsequently infected with SARS-CoV-2-S^D614G^ (green) K18-hACE2 mice were challenged intranasally with 3x 10^4^ TCID_50_ SARS-CoV-2-S^D614G^. (**a**) Determination of virus neutralization titers (ND_50_) in serum of offspring 1 week before and 5 days after challenge infection. (**b**) Body weight at the indicated times pi. (**c**) Determination of infectious SARS-CoV-2 titers in lung and brain tissue homogenates 5 days pi. Geometric means and geometric SD are indicated. Statistically significant differences were determined by the two-way ANOVA test (**a**) or by the two-tailed unpaired Student’s t test (**c**) (*p < 0.05; **p < 0.005, ****p < 0.0001).

### The VSV*ΔG-S_Δ21_ vaccine candidate induces broadly reactive neutralizing antibodies

In the course of the ongoing COVID-19 pandemic, a number of SARS-CoV-2 variants have emerged that revealed characteristic mutations in the spike protein that improved virus fitness and/or allowed escape from antibody-mediated virus neutralization.^19^ To determine how broadly the neutralizing antibodies induced by the three VSV-vectored vaccines would react, we performed a pseudotype virus neutralization assay using VSV*ΔG(Fluc) replicon particles that were *trans*-complemented with the S protein derived either from the Hu-1 reference strain (harboring the D614G mutation), or from the variants of concern (VOCs) Alpha, Beta, Gamma, and Delta. Mouse immune sera collected 4 weeks after boost with VSV*ΔG-S_Δ21_ neutralized pseudotype virus bearing either the S^Alpha^ or S^Beta^ equally well than pseudotype virus bearing the S^D614G^ protein (**Fig. 6a**). In contrast, neutralization of pseudotype viruses harboring S^Gamma^ and S^Delta^ was significantly reduced (**Fig. 6a**). Sera from mice vaccinated with the VSV*M_q_ΔG-S_Δ21_ vaccine equally neutralized S^D614G^ and S^Alpha^ pseudotype viruses, but were significantly less efficient against S^Beta^, S^Gamma^, and S^Delta^ pseudotype viruses (**Fig. 6b**). Finally, serum antibodies from mice vaccinated with the VSV*ΔG-G_RBD_ vaccine showed reduced neutralizing activity against the four different VOC spike pseudotype viruses (**Fig. 6c**). Together, these results indicate that among the three vaccine candidates tested, the VSV*ΔG-S_Δ21_ vaccine induced the broadest humoral immune response.

**Fig. 6.**
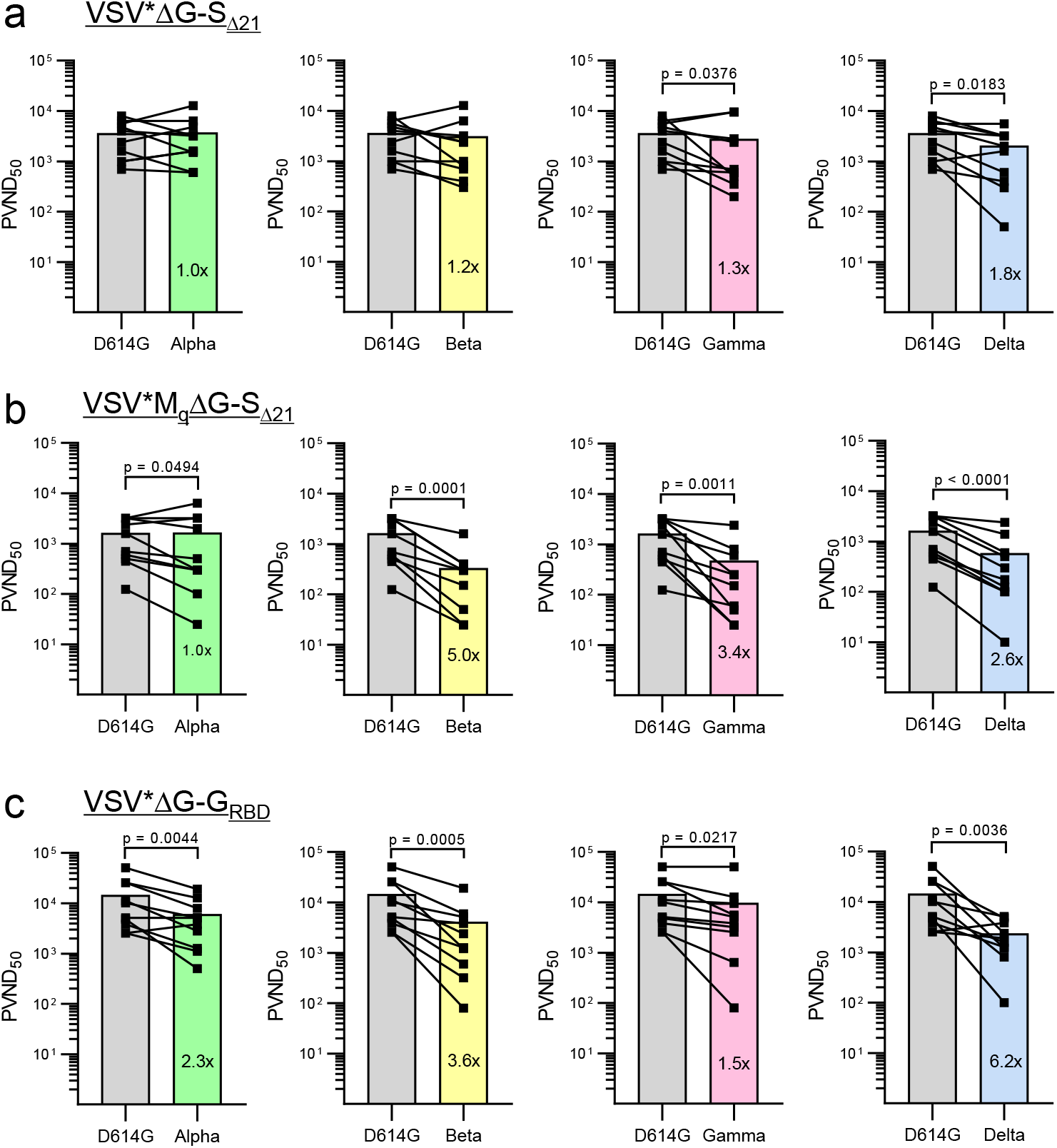
Neutralizing antibodies induced by VSV-vectored vaccines differ in their capacity to block virus entry mediated by spike proteins derived from SARS-CoV-2 variants. Immune sera were prepared from K18-hACE2 mice 4 weeks after the second immunization with either (**a**) VSV*ΔG-S_Δ21_ (n = 10), (**b**) VSV*M_q_ΔG-S_Δ21_ (n = 10), and (**c**) VSV*ΔG-S_RBD_ (n = 5). Virus neutralization tests were performed using the VSV*ΔG(FLuc) vector pseudotyped with the S protein of the indicated SARS-CoV-2 VOC. At 20h pi of Vero E6 cells, the reciprocal serum dilution causing 50% inhibition of the pseudotype virus-encoded firefly luciferase reporter expression was determined and expressed as pseudotype virus neutralization dose 50% (PVND_50_). Lines connect the corresponding PVND_50_ titers of individual immune sera. The bars represent the mean neutralization titers. Statistically significant differences as determined by the two-tailed paired Student’s t test are indicated.

### The VSV*ΔG-S_Δ21_ vaccine protects K18-hACE2 mice less efficiently from infection with the SARS-CoV-2 Delta variant

To determine whether immunization with VSV*ΔG-S_Δ21_ also protects from a heterologous challenge *in vivo*, K18-hACE2 mice were firstly immunized twice with 2×10^6^ ffu of VSV*ΔG-S_Δ21_. Vaccination induced high VSV*ΔG-S_Δ21_ neutralizing activity, which was increased after boost (**Fig. 7a**). Sera of boosted mice had reduced neutralizing activity against pseudotype virus *trans*-complemented with S^Delta^ than against the S^D614G^ pseudotype (**Fig. 7b**). Immunized and naïve mice were infected with SARS-CoV-2^Delta^ VOC via the nasal route using a dose of 5×10^4^ pfu per animal and followed up to day 5 pi. Weight loss was observed in 4 out of 5 naïve mice and in 1 out of 5 vaccinated mice (**Fig. 7c**). The four naïve mice showing weight loss, but none of the vaccinated mice, presented typical clinical signs of central nervous system infection (hunched posture, ruffled fur, tremors, or ataxia) on day 5 pi and were euthanized as planned. Virus loads measured by qPCR were significantly reduced in oropharyngeal swabs and organs collected from the immunized animals at day 5 pi compared naïve mice (**Fig. 7d, e**). Critically, no SARS-CoV-2^Delta^ plaques were grown from lungs and brains of all vaccinated mice, whereas all control mice had high virus titers on day 5 pi (**Fig. 7f**). Surprisingly, lung pathology scores of the immunized and naïve groups did not significantly differ from each other, even in the absence of detectable viral antigen by IHC in the vaccinated mice (**Supplementary Fig. 4a**). Another five vaccinated K18-hACE2 mice were surveyed up to 14 days following infection with SARS-CoV-2^Delta^. While two animals stayed healthy during this time period, three mice showed significant weight loss of which one reached exclusion criteria at day 5 pi, while the other two mice recovered until day 14 pi (**Fig. 7g**). Analysis of the serum of the surviving animals revealed significantly increased neutralizing antibody titers compared to the pre-challenge (boost) sera (**Fig. 7h**), indicating that the humoral immune response to the VSV-vectored spike antigen was further boosted by challenge infection with SARS-CoV-2^Delta^. Together, these data indicate that intramuscular immunization of K18-hACE2 mice with the VSV*ΔG-S_Δ21_ vector provides partial protection against SARS-CoV-2^Delta^ VOC, reducing virus load in the respiratory tract, blocking dissemination to the brain, but without preventing lung pathology.

**Fig. 7.**
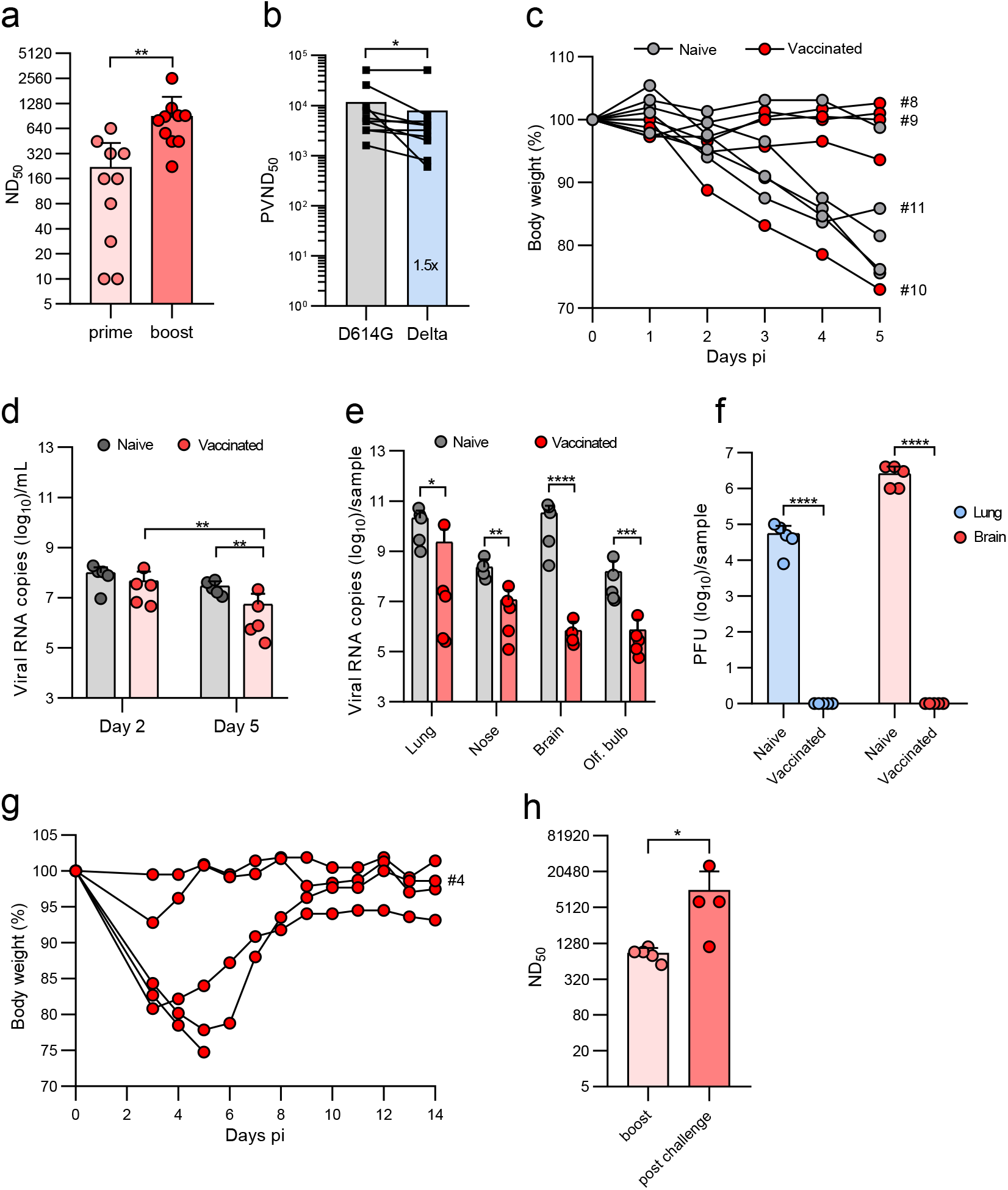
The VSV*ΔG-S_Δ21_ vaccine partially protects K18-hACE2 mice from SARS-CoV-2^Delta^ infection. K18-hACE2 mice (n = 10) were immunized (i.m.) twice with 10^6^ ffu of VSV*ΔG-S_Δ21._ (**a**) Determination of VSV*ΔG-S_Δ21_-neutralizing antibodies in serum collected 3 weeks after the first (prime) and 3 weeks after the second immunization (boost). (**b**) Determination of the pseudotype virus neutralization dose 50 (PVND_50_) using the VSV*ΔG(FLuc) reporter virus pseudotyped with the S^D614G^ or S^Delta^ protein. (**c-h**) At 4 weeks after the second immunization, animals were infected with 8×10^4^ pfu of SARS-CoV-2^Delta^ VOC via the nasal route. A group of naïve K18-hACE2 mice (n = 5) served as control. (**c**) Relative body weight of mice in the course of infection up to day 5 pi. Some Individual curves were labeled with an ID number to identify the animals in the histological analysis (**Supplementary Fig. 4A**). (**d, e**) Quantification of SARS-CoV-2 RNA copies by RT-qPCR in oropharyngeal swab samples collected at days 2 and 5 pi (**d**), and in the indicated organs prepared at day 5 and 14 pi (**e**). (**f**) Determination of infectious SARS-CoV-2^Delta^ titers in homogenates of lung and brain at day 5 pi. (**g**) Relative body weight of vaccinated mice in the course of infection with SARS-CoV-2^Delta^ up to day 14 pi. (**h**) Determination of neutralizing serum antibody titers (ND50) 18 days after the boost and 14 days post challenge infection. Mean values and SD are shown. Statistical significance was calculated by the unpaired Student’s t test (**a, e, f, h**), by the paired Student’s t test (**b**), and by the two-way ANOVA test (**d**) (*p < 0.05, **p < 0.01; ***p < 0.005; ****p < 0.0001).

## Discussion

Recently, a propagation-competent VSV-vectored vaccine that encode a SARS-CoV-2 spike protein with a truncated cytoplasmic tail was shown to induce protective immune responses against SARS-CoV-2 in hamster and mouse models.^4, 5, 15^ However, a similar vaccine candidate (V590) generated by Merck did not induce a sufficiently high immune response when evaluated in a phase I trial involving 252 participants and was therefore discontinued (www.clinicaltrials.gov/ct2/show/NCT04569786). In the present work, we followed up on these studies and evaluated an analogous VSV-vectored vaccine, VSV*ΔG-S_Δ21_. Replication of this chimeric virus fully relied on the recombinant SARS-CoV-2 spike protein, which mediates ACE2-dependent infection. However, intramuscular immunization of K18-hACE2 and C57BL/6J mice with non-trans-complemented VSV*ΔG-S_Δ21_ triggered the production of spike-specific antibodies at only very low levels, suggesting that skeletal muscle cells at the vaccine inoculation site were resistant to infection with VSV*ΔG-S_Δ21_ due to low-level or absent ACE2 expression.

When VSV*ΔG-S_Δ21_ was propagated on helper cells providing the VSV G protein *in trans*, both glycoproteins were incorporated into the viral envelope (**Fig. 1c and d**). Thus, this helper cell-grown chimeric virus can infect cells in an ACE2-independent manner. The VSV G protein is known to mediate infection of a very broad range of cell types from different species,^20^ such as skeletal muscle cells, and thus enhanced immunization efficacy by VSV G protein *trans*-complemented VSV*ΔG-S_Δ21_. Likewise, immunization with the helper cell-grown VSV*ΔG-S_Δ21_ induced neutralizing antibodies at high levels and protected K18-hACE2 mice from challenge infection with SARS-CoV-2.

The present study demonstrates that intramuscular immunization with helper cell-grown VSV*ΔG-S_Δ21_ completely protected K18-hACE2 mice from lethal SARS-CoV-2-S^D614G^ challenge infection. Indeed, dissemination of the challenge virus to the brain, which is known to cause rapid death of K18-hACE2 mice,^18^ was prevented in the vaccinated animals. Moreover, virus load in the lung of vaccinated animals was largely reduced, although some infectious foci were still detectable by IHC analysis in lung sections 5 days pi. Likewise, vaccinated mice still revealed some pulmonary changes at 14 days pi, when the virus was mostly cleared. It remains unclear whether these changes are due to virus-mediated damage to the lung or a consequence of the immune response. Collectively, our data support the hypothesis that intramuscular immunization with VSV*ΔG-S_Δ21_ induced spike-specific serum antibodies, which inhibited systemic dissemination of SARS-CoV-2 to extrapulmonary organs such as the brain. A fraction of serum IgG may have been transported across epithelial barriers and secreted into the lower respiratory tract,^21^ where these antibodies could interfere with virus infection and dissemination. However, the availability of serum antibodies in the respiratory tract might not be sufficient to completely prevent SARS-CoV-2 replication in the airways.

Compared to VSV*ΔG-S_Δ21_, the VSV*M_q_ΔG-S_Δ21_ vector vaccine induced lower levels of neutralizing antibodies, which were less effective against some SARS-CoV-2 VOCs. As VSV*M_q_ΔG-S_Δ21_ vector encoded the same antigen as VSV*ΔG-S_Δ21_, the mutant M_q_ gene is likely responsible for the 2-5 fold reduction in heterologous neutralization performance against VOCs. As the modified matrix protein M_q_ lacks host shut-off activity, VSV*M_q_ΔG-S_Δ21_ is less cytotoxic than VSV*ΔG-S_Δ21_,^16^ which means that there might be less danger-associated molecular patterns (DAMPs) released from infected cells that can stimulate the immune system. Concomitantly, VSV*M_q_ΔG-S_Δ21_ was unable to block the synthesis and secretion of type I interferon (IFN), which likely led to reduced viral replication, transcription and protein synthesis in vector-infected cells.^20^ Consequently, lower amounts of vector-encoded vaccine antigen might have been produced, potentially explaining why lower levels of virus-neutralizing antibodies were elicited by this vector.

In addition to the propagation-competent VSV*ΔG-S_Δ21_ and the propagation-restricted VSV*M_q_ΔG-S_Δ21_, we produced a fully propagation-defective vector vaccine, which encoded a chimeric membrane protein, G_RBD_, integrating parts of the VSV G protein with the RBD of the SARS-CoV-2 spike protein. As this viral envelope protein was devoid of any domains involved in membrane fusion, the VSV*ΔG-G_RBD_ vector could only be propagated on helper cells providing the VSV glycoprotein *in trans*. Although this vector encoded the wild-type M protein, it was less immunogenic than VSV*ΔG-S_Δ21_ and a second immunization was required to reach appreciable levels of neutralizing antibodies in K18-hACE2 mice. The reduced immunogenicity of this vector vaccine might be due to an inferior RBD conformation or presentation by the synthetic membrane protein. Nevertheless, the immune response induced after two immunizations with VSV*ΔG-G_RBD_ was sufficient to protect K18-hACE2 mice from challenge with a lethal dose of SARS-CoV-2. Interestingly, a very similar RBD-based VSV-vectored vaccine has recently been evaluated and was found to induce a protective immune response in K18-hACE2 mice after a single intramuscular immunization.^22^

Since the onset of the pandemic in December 2019, several SARS-CoV-2 VOCs have emerged that revealed an altered transmissibility/infectivity and frequently showed mutations in the RBD and/or N-terminal domain (NTD) of the spike protein, which allowed the virus to escape neutralizing antibodies.^23–25^ In Europe, the original SARS-CoV-2 was initially replaced by the Alpha, followed by the Delta and recently by the Omicron variant.^26^ Preferentially, a vaccine should induce a broadly reactive immune response by targeting epitopes in the spike protein that are conserved among different VOCs. When immune sera of vaccinated mice were tested for neutralizing antibodies against the different VOCs taking advantage of pseudotype viruses, antibodies induced by the VSV*ΔG-S_Δ21_ vaccine inhibited virus entry mediated by the VOC spike proteins fairly well with less than a 2-fold reduction in neutralizing titers. In contrast, immune sera from mice that had been vaccinated with the VSV*ΔG-G_RBD_ vector revealed a less broadly neutralizing activity, in particular against the Delta spike protein. As this vaccine candidate only presents the RBD, it might be easier for new SARS-CoV-2 variants to escape neutralizing antibodies.

K18-hACE2 mice vaccinated with VSV*ΔG-S_Δ21_ were fully protected against SARS-CoV-2^Delta^ VOC dissemination to the brain and no virus titers were detectable in lungs and brain. However, vaccinated and naïve mice showed similar lung pathology at day 5 and day 14 pi suggesting that early virus replication occurred also in the lungs of vaccinated animals with consequent tissue damage. Moreover, in four out of ten vaccinated mice challenge infection resulted in clinical disease and body weight loss. These findings suggest that both antigen mismatch and the low availability of serum antibodies in lung tissues following intramuscular immunization may explain the reduced protection of K18-hACE2 mice from breakthrough infection with the Delta variant.

Since the onset of the pandemic, several vaccine platforms for protection against SARS-CoV-2 have been developed. In particular, mRNA-based COVID-19 vaccines have been rapidly developed and licensed, and are now the most widely used vaccines for SARS-CoV-2 immunoprophylaxis.^27, 28^ However, mRNA-based vaccines require at least two intramuscular immunizations to achieve a fully protective immune response to SARS-CoV-2.^29, 30^ In this regard, VSV-vectored spike protein may offer a clear advantage over mRNA-based vaccines because they rapidly induce a protective immune response already after a single immunization.^5, 31, 32^ In line with these findings, our study demonstrates that a single intramuscular immunization with VSV*ΔG-S_Δ21_ or VSV*M_q_ΔG-S_Δ21_ induces neutralizing serum antibody at levels that can mediate protection as evident from challenge infection of mouse pups that had received maternal antibodies. One reason for the superior immunogenicity of VSV-vectored vaccines is that in contrast to mRNA vaccines the viral RNA vector genome is amplified in the cytosol of the host cell, which leads to extremely high antigen expression levels and strong stimulation of the immune system. We therefore anticipate that vaccinated humans would develop a protective immune response shortly after one intramuscular immunization. However, a second vaccination is still recommended as it will further improve the availability of spike-specific antibodies in the airways and further limit virus replication and lung pathology.^4^ High titers of neutralizing antibodies after a second immunization will also help to better protect from replication of emerging VOCs as these variants are neutralized with significantly lower efficacy.

Our data also suggest that intramuscularly applied COVID-19 vaccines may not sufficiently prevent SARS-CoV-2 replication in the respiratory tract, which could lead to still severe lung pathology. This problem may become even more difficult when neutralizing antibody titers drop with time after the immunization, or if the vaccinated person encounters a SARS-CoV-2 variant harboring escape mutations in the spike. Furthermore, intramuscular immunization may not efficiently prevent virus transmission, which would be favorable with respect to control the pandemic. In order to efficiently block the chain of airborne infection with SARS-CoV-2, vaccines may be applied via the mucosal rather than the intramuscular route. Immunization via the intramuscular route can induce antibodies of the IgA isotype, which are efficiently secreted into the mucosal tissues of the respiratory tract where they can directly neutralize pathogens at the site of primary replication.^33, 34^ In contrast to the presently approved mRNA vaccines, which rely on intramuscular administration, there is already first evidence from the hamster model that mucosally applied VSV-vectored COVID-19 vaccines can in principle be very effective in preventing virus shedding from the respiratory tract.^32, 35^ Recently, VSV-vectored spike antigen that was administered via the oral route efficiently boosted the immune response to SARS-CoV-2 in nonhuman primates.^36^

In summary, our findings demonstrate that *trans*-complementation of VSV-vectored vaccines with the VSV G protein has multiple impacts: (1) it improves virus yield to maximize vaccine production, (2) it allows efficient ACE2-independent intramuscular immunization leading to dramatically enhanced immune responses even after a single administration, (3) it supports the induction of broadly neutralizing antibodies when associated with the proper antigen. Future development of VSV-vectored vaccines will include improved induction of protective immune responses against conserved epitopes and enhanced mucosal immunity.

## Materials and Methods

### Cells

Vero E6 cells were kindly provided by Christian Drosten/Marcel Müller (Charite, Berlin, Germany) and maintained in Dulbecco’s minimal essential medium (DMEM; Life Technologies) supplemented with 10% fetal bovine serum (FBS) and non-essential amino acids (Life Technologies). Vero cells expressing human TMPRSS2 (Vero-hTMPRSS2) were kindly provided by Markus Hoffmann (German Primate Center, Göttingen, Germany) and maintained DMEM supplemented with 10% FBS, non-essential amino acids and the selection antibiotic puromycin (0.5 μg/ml). A549 human lung carcinoma cells were obtained from the German cell culture collection DSMZ (Braunschweig, Germany) and grown in Ham’s F12/DMEM medium with 10% FBS. A549 cells stably expressing human ACE2 and TMPRSS2 were obtained from Invivogen (Toulouse, France) and maintained in DMEM medium with 10% FBS and selection antibiotics puromycin (0.5 μg/ml) and hygromycin B (300 μg/ml). BHK-21 cells were obtained from American Type Culture Collection (ATCC® CCL-10; Manassas, VA, USA) and maintained in Glasgow’s minimal essential medium (GMEM, Life Technologies) supplemented with 5% FBS. BHK-G43, a transgenic BHK-21 cell clone expressing the VSV G protein in a regulated manner,^17^ was maintained in GMEM containing 5% FBS. I1-Hybridoma were purchased from ATCC® (CRL-2700™) and maintained in minimal essential medium (MEM, Life Technologies) supplemented with 15% FBS.

### Viruses

SARS-CoV-2-S^D614G^ encoding the spike D614G mutation was generated by reverse genetics using a novel synthetic genomics platform.^37^ The SARS-CoV-2 Delta variant B.1.617.2 used in this study was isolated from COVID-19 patient in India. The genomic sequence of this virus has been completely deciphered (GISAID accession number EPI_ISL_2535433).^38^ VSV*ΔG is a propagation-deficient VSV which lacks the glycoprotein G gene but encodes the GFP reporter protein.^16^ VSV*ΔG(FLuc) is a glycoprotein-deficient VSV encoding GFP and firefly luciferase.^39^ VSV*ΔG and VSV*ΔG(FLuc) were propagated on transgenic BHK-G43 cells expressing the VSV G protein in a regulated manner.^17^ The *trans*-complemented particles were stored at −70°C in the presence of 5% FBS. Pseudotype viruses were titrated on BHK-21 cells that were grown in 96-well cell culture plates.^39^

### Plasmids

SARS-CoV-2 harboring the D614G mutation in the spike protein was isolated from a COVID-19 patient at the University hospital in Freiburg (Germany) in spring 2020. The spike cDNA (GISAID accession number: EPI_ISL_852667) was amplified without the region encoding the 18 C-terminal amino acids and cloned into the pCAGGS expression plasmid taking advantage of KpnI and XhoI endonuclease restriction sites. The expression plasmids pLV-spikev2, pLV-spikev3, and pLV-spikev5 encoding the spike proteins of the Alpha variant (B.1.1.7) (GISAID accession number: EPI_ISL_811870), the Beta variant (B.1.351) (GISAID accession number: EPI_ISL_745146, and the Gamma variant (P.1) (GISAID accession number: EPI_ISL_811149), respectively were purchased from Invivogen (Toulouse, France). All three encoded spike variants were truncated at the C terminus by 19 amino acids. The pCDNA3.3-SARS2-B.1.617.2 plasmid encoding the spike protein of the Delta variant (B.1.617.2) (GenBank accession number: 43740568) has been published by David Namazee (Scripps Research Institute, La Jolla, CA, USA)^40^ and was obtained via Addgene (Cat. No. 172320). The C-terminus of the spike protein encoded by this plasmid was truncated by 18 amino acids.

### Generation of recombinant VSV vector vaccines

Recently, we generated a recombinant chimeric virus, VSV*ΔG(MERS-S), in which the VSV glycoprotein (G) gene was replaced by the full-length MERS-CoV spike protein.^41^ VSV*ΔG(MERS-S) also encoded a GFP reporter which was expressed from an additional transcription unit located downstream of the spike gene. To generate a chimeric VSV expressing the SARS-CoV-2 spike protein, the MERS-S gene in the antigenomic plasmid pVSV*ΔG(MERS-S) was replaced by a modified SARS-CoV-2 spike gene (Genscript, Piscattaway, USA) taking advantage of the flanking MluI and BstEII endonuclease restriction sites. The modified SARS-CoV-2 spike gene was based on the Wuhan-Hu-1 strain (GenBank accession no.: NC_045512) but lacked the region encoding the C-terminal 21 amino acids and contained the mutations R685G, H655Y, D253N, W64R, G261R, A372T, and R685G.^15^

The VSV matrix protein quadruple mutant M_q_ has been shown to lack host shut-off activity.^16^ To generate a VSV-vectored vaccine candidate with reduced cytotoxicity, the pVSV*ΔG-S_Δ21_ genomic plasmid was modified by replacing the XbaI/MluI fragment (including the M gene) with the corresponding fragment of the pVSV*M_q_ plasmid^16^, resulting in the plasmid pVSV*M_q_ΔG-S_Δ21_.

To generate a propagation-defective VSV vaccine vector which displays the receptor-binding domain (RBD) of the SARS-CoV-2 spike protein, a synthetic cDNA encoding the N-terminal 23 amino acids of the VSV G protein (GenBank accession no.: J02428) including the signal peptide (aa 1-16), the RBD of the SARS-CoV-2 (Wuhan-Hu-1 strain) spike protein (aa 323-550), and the C-terminal region of the VSV G protein (aa 421-511) comprising the stem region, the transmembrane domain and the cytoplasmic domain. was synthesized by Genscript Inc. (Leiden, The Netherlands) and inserted into the pVSV*ΔG(MERS-S) plasmid thereby replacing the MERS-CoV S gene. The resulting plasmid was designated pVSV*ΔG-G_RBD_.

The recombinant viruses were rescued from the cell culture supernatant of BHK-G43 cells following infection of the cells with a modified vaccinia virus Ankara encoding T7 RNA polymerase and transfection with a set of four plasmids, one encoding the recombinant VSV genome and the others encoding the VSV N, P, and L genes, all under control of the T7 promotor.^42^ At 48 hours post transfection, the cell culture supernatant was clarified by low-speed centrifugation and passed through a 0.2-µm pore size filter for depletion of the vaccinia virus. The recombinant viruses were propagated on Vero E6 cells up to passage 4. The recombinant viruses were stored in aliquots at −70°C in the presence of 10% FBS. Infectious virus titers were determined on confluent Vero E6 cells grown in 96-well microtiter plates. The cells were inoculated in duplicate with 40 µl per well of serial 10-fold virus dilutions for 1 hour at 37°C. Thereafter, 160 µl of EMEM containing 1% methyl-cellulose was added to each well, and the cells were incubated for 24 hours at 37°C. The number of infectious foci was determined under the fluorescence microscope taking advantage of the GFP reporter and infectious virus titers were calculated and expressed as ffu/ml.

### Enzyme-linked immunosorbent assay

The prefusion-stabilized ectodomain of the SARS-CoV-2 spike protein containing a C-terminal trimerization motif, an HRV 3C protease cleavage site, and a twin strep-tag was expressed and purified as recently described. ^43, 44^ The antigen (1 µg protein/100 µl/well) was coated overnight at 4°C onto MaxiSorp 96-well ELISA plates (Nunc). Any remaining protein binding sites on the polystyrene surface were subsequently blocked by adding 250 µl/well of PBS containing 1% (w/v) of bovine serum albumin (BSA, Sigma). The plates were incubated overnight at 4°C followed by three wash steps with PBS containing 0.01% (v/v) Tween 20. The immune sera were serially diluted in PBS containing 0.1% BSA and incubated for 60 minutes at room temperature with the immobilized spike antigen. Thereafter, the ELISA plates were washed four times with PBS/0.01% Tween 20 and subsequently incubated for 60 minutes at room temperature with goat anti-mouse immunoglobulins conjugated to horseradish-peroxidase (Agilent-Dako, Cat. No.: P0260, 1:5000 in PBS/0.1%BSA). The ELISA plates were washed 5 times with PBS/0.01% Tween 20 and then incubated for 10 minutes at room temperature with 50 µl/well of 3,3’5,5’-tetramethylbenzidine (TMB) peroxidase substrate (Sigma). After the reaction was stopped by adding 50 µl/well of HCl (1 M), the optical density (OD) at 450 nm was determined with a GloMax plate reader (Promega). The cutoff OD_450_ value for antigen-specific binding was set as the mean OD_450_ caused by mouse pre-immune serum plus 0.2 OD_450_ units (corresponding approximately to a tenfold standard deviation of the blank). The antibody ELISA titer was defined as the reciprocal value of the highest serum dilution causing an OD_450_ signal above the cutoff value.

### Neutralization assay using propagation-competent chimeric VSV*ΔG-S_Δ21_

Twofold serial dilutions of heat-inactivated immune sera/plasma were prepared in quadruplicates in 96-well cell culture plates using DMEM cell culture medium (50 µl/well). To each well, 50 µl of DMEM containing 100 ffu of VSV*ΔG-S_Δ21_ were added and incubated for 60 min at 37°C. Subsequently, the antibody/virus mix was transferred to confluent Vero E6 cell monolayers in 96-well cell culture plates and incubated at 37°C for 24 hours. Infected cells were detected by fluorescence microscopy taking advantage of the virus-encoded GFP reporter. A single infection event per well was scored as non-protected. Neutralization doses 50% (ND_50_) values were calculated according to the Spearman and Kärber method.^45^

### Neutralization assay using VSV*ΔG(FLuc) pseudotypes

To test immune sera for neutralizing activity against SARS-CoV-2 spike variants, single cycle VSV*ΔG(FLuc) reporter virus was pseudotyped with either the D614G spike protein mutant or with the spike proteins of one of the variants Alpha (B.1.1.7), Beta (B.1.351), Gamma (P.1) and Delta (B.1.617.2) according to a recently described procedure.^46^ Briefly, the pseudotype viruses were produced by transfecting BHK-21 cells with the respective spike expression plasmid using Lipofectamine 2000 (Life Technologies, Zug, Switzerland) as transfection reagent. At 16 hours pi, the cells were washed one with GMEM and incubated for 4 hours with GMEM with 5% FBS. The transfected cells were subsequently inoculated for 60 min at 37°C with VSV*ΔG(FLuc) using an moi of 5 ffu/cell. The cells were washed twice with GMEM and subsequently incubated at 37°C with GMEM containing 5% FBS and 10% conditioned cell culture medium from hybridoma cells secreting the neutralizing monoclonal antibody Mab I1. At 20 hours pi, the cell culture supernatant was harvested, cell debris removed by low-speed centrifugation. The pseudotype virus suspension was buffered by addition of 50 mM HEPES (Gibco) and stored in aliquots at −70°C. The pseudotype viruses were titrated on Vero E6 cells using the GFP reporter for detection of infected cells.

The pseudotype virus neutralisation (PVN) test was performed by incubating twofold serial dilutions of the murine immune sera (heat-inactivated at 56°C, 30 minutes) with pseudotype virus (50 μl/well containing 200 ffu) for 60 min at 37°C. The serum/virus mix was incubated for 16 hours at 37°C with confluent Vero E6 cells in 96-well cell culture plates. The cells were lysed by adding 100 µl of luciferase lysis buffer (Biotium, Fremont, CA, USA). The lysates (50 µl) were transferred to white microtiter plates and 50 µl of firefly luciferase ONE-Glo^TM^ substrate (Promega, Dübendorf, Switzerland) added. Luminescence was recorded using a GloMax® plate reader (Promega). The reciprocal antibody dilution causing 50% reduction of the luminescence signal was calculated and expressed as pseudotype virus neutralization dose 50% (PVND_50_).

### Interferon bioassay

A549-hACE2-hTMPRSS2 cells were infected with either VSV*ΔG-S_Δ21_ or VSV*M_q_ΔG-S_Δ21_ using an moi of 1 ffu/cell and aliquots of the conditioned cell culture supernatant were collected at 24 and 48 hours pi. Antiviral activity in the cell culture supernatants was determined on normal A549 cells using an established bioassay taking advantage of VSV*ΔG(Fluc) as reporter virus.^39^ Briefly, A549 cells grown in 96-well cell culture plates were incubated for 20 hours at 37°C with serial twofold dilutions of the conditioned supernatant and subsequently inoculated for 5 hours at 37°C with VSV*ΔG(Fluc) using an moi of 5 ffu/cell. The cells were lysed and luciferase activity monitored in a GloMax® plate reader (Promega, Dübendorf, Switzerland) using the One-Glo^TM^ luciferase substrate (Promega). The reciprocal dilution of the cell culture supernatant leading to 50% reduction of firefly luciferase expression was calculated (inhibitory concentration 50%, IC_50_).

### Western blot

Confluent monolayers of BHK-G43 cells grown in T75 flasks (about 10^7^ cells) were treated for 6 hours with mifepristone (10^-9^ M) and subsequently infected with either VSV*ΔG-S_Δ21_, VSV*M_q_ΔG-S_Δ21 or_ or VSV*ΔG-G_RBD_ using an moi of 0.1 ffu/cell. In parallel, Vero E6 cells that were grown in T75 flasks (about 10^7^ cells/flask) were infected with either VSV*ΔG-S_Δ21_ or VSV*M_q_ΔG-S_Δ21_ using an moi of 0.1 ffu/cell. At 24 hours post infection, the cell culture supernatant of the infected cells were collected and cell debris removed by centrifugation (1200 x g, 15 min, 4°C). Subsequently, virus particles were pelleted from the clarified cell culture supernatant by ultracentrifugation through a 25% (w/w) sucrose cushion and directly solubilized by adding preheated (95°C) sodium dodecyl sulfate (SDS) sample buffer with or without 0.1 M of dithiothreitol (DTT) to the pellets. The solubilized viral proteins were subsequently separated by SDS polyacrylamide gel electrophoresis (PAGE) using 4 - 12% gradient gels (<u>SurePAGE™</u>; Genscript, Leiden, The Netherlands) and transferred to nitrocellulose membranes by semi-dry blotting. The nitrocellulose membrane was blocked overnight at 4°C with Odyssey Blocking Reagent (Li-COR Biosciences, Lincoln, NE) and subsequently incubated with either reconvalescent COVID-19 immune serum (ND_50_ of 960),^46^ diluted 1:1000 in PBS or a polyclonal rabbit immune serum which recognizes the VSV G and M antigens.^16^ The membrane was washed four times with PBS containing 0.1% Tween-20 and incubated with the secondary antibodies IRDye 800CW goat anti-human IgG (LI-COR Biosciences, Cat. no. 926-322232), and IRDye 680RD goat anti-rabbit IgG (LI-COR Biosciences, Cat. no. 926-68071), both diluted 1:5000 in PBS. Following several washing steps with PBS/0.1% Tween 20 the blots were scanned with the Odyssey Infrared Imaging system (LI-COR Biosciences, Bad Homburg, Germany). The unprocessed Western blot results are shown in **Supplementary Figure 5**.

### Animal experiments

Animal experiments have been performed in compliance with the Swiss animal protection law and approved by the animal welfare committee of the Canton of Bern (authorization number BE43/20). Based on previous experiments, we aimed to detect a 10-fold reduction in virus load between two groups with a standard deviation equal to the mean value of the vaccinated group. Power calculations assuming an alpha error of 0.05 and a beta error of 0.80 indicated that statistical significance will be reached in groups of 5 mice. Calculations were made using the online software of Columbia University Medical Center (www.biomath.info).

Male, 12-18 week-old, C57BL/6 or female 17-41 week-old hemizygous transgenic mice (Tg(K18-hACE2)2Prlmn; referred to as K18-hACE2)^47–49^ were bred at the specific pathogen-free facility of the Institute of Virology and Immunology (Mittelhäusern, Switzerland). The mice were kept in individually ventilated cages (blue line, Tecniplast) under the following conditions: 12-h/12-h light/dark cycle, 22 ± 1 °C ambient temperature and 50 ± 5% humidity, autoclaved food and acidified water. Mice were transferred to individually HEPA-filtered cages (IsoCage N, Tecniplast) in age- and sex-matched groups and immunized intramuscularly by injection of 20-25 μl of GMEM containing 10^5^ or 10^4^ ffu of Vero E6 or BHK-G43 cell-grown VSV*ΔG-S_Δ21_. After 4 weeks, the animals were immunized a second time using the same vector vaccine, route and dose. Blood (approximately 200 µl/animal) was collected from the animals via the retrobulbar route under brief isoflurane anesthesia one week before and three weeks after the boost. Sera were prepared by centrifugation of coagulated blood and stored in aliquots at −20°C. Four weeks after boost immunization, the K18-hACE2 mice were transferred to the biosafety level 3 (BSL-3) facility of the Institute of Virology and Immunology, briefly anesthetized with isoflurane and challenged intranasally with 10^5^ pfu of SARS-CoV-2-S^D614G^ (20 μl per nostril). Infected mice were monitored daily for activity, posture and hair grooming. Body weight was measured daily and changes are presented as percent of body weight at time of infection. Humane endpoint for euthanasia were reached when one or more of the following symptoms are observed: Body weight loss >25%, significant dehydration, ataxia, or circling. Oropharyngeal swabs were collected under brief isoflurane anesthesia using ultrafine sterile flock swabs (Hydraflock, Puritan, 25-3318-H). Five days after challenge, the mice were euthanized and organs were sampled for histological analysis, virus titration, and RT-qPCR.^37^

To compare three different vector vaccine candidates, ten- to 22-week-old female K18-hACE2 mice and six-week-old male were immunized twice via the intramuscular route 4 weeks apart with 2 × 10^6^ ffu of the respective VSV vector suspended in 10 μl of PBS. Age- and sex-matched groups were used for each round of immunization/challenge. On days 25 and 55 after the initial immunization, blood was collected and serum samples were stored at −20°C for further analysis. Mice were infected intranasally as described above. Mice were sacrificed once reached the humane end-point or at 5 and 14 days pi.

To analyze vaccine protection against the SARS-CoV-2^Delta^ variant, 14-40-week-old K18-hACE2 mice were immunized and boosted with the 10^6^ ffu of BHK-G43 cell-grown VSV*ΔG-S_Δ21_ and challenged as described above, with 8×10^4^ pfu of SARS-CoV-2^Delta^ isolate. Five naïve, 7-week-old female K18-hACE2 were used as control for this experiment. Mice were sacrificed once reached the humane end-point or at 5 and 14 days pi.

### Flow cytometry and tetramer staining

Alexa Fluor 647-labelled SARS-CoV-2 S 539-546 tetramers preloaded with VNFNFNGL peptide (S-tet) were provided by the National Institutes of Health tetramer core facility. The fluorescence-minus-one control and staining with the irrelevant influenza A NP 366-374 tetramer were performed to set the gates. EDTA-anticoagulated whole blood (50 µL) were stained with PerCP/Cyanine5.5 anti-mouse CD45 antibody, clone 30-F11 (Biolegend, Cat. no. 103132), PE anti-mouse CD3epsilon antibody, clone 145-2C11 (Biolegend, Cat. no. 100308), FITC anti-mouse CD8a antibody, clone 53-6.7 (Biolegend, Cat No. 100706). Dead cells were excluded with LIVE/DEAD fixable Aqua Dead staining kit (ThermoFisher Scientific, L34957). Blocking was done with CD16/CD32 (clone 2.4G2, Biolegend) and avidin (Millipore) prior to staining with cell surface antibodies and tetramer. Acquisition of data was performed on a FACS-Canto II (BD Biosciences) and analyzed with FlowJo software version 10.

### RT-qPCR

To determine the virus load in oropharyngeal swab samples, the swab tips were placed in 0.5 ml of RA1 lysis buffer (Macherey-Nagel, ref. 740961) containing 1% (v/v) of β-mercaptoethanol. After a short centrifugation step, RNA was extracted from 200 μl of the lysate using the Nucleo-Mag Vet kit (Macherey Nagel). To detect viral RNA in lungs and brains of infected mice, the superior right lobe of the lung and the frontal half of the right brain hemisphere were homogenized in 0.5 ml of RA1 lysis buffer supplemented with 1% β-mercaptoethanol using a Bullet Blender Tissue Homogenizer (Next-Advance Inc.) The homogenate was clarified by centrifugation and RNA extracted from 100 μl of the cleared supernatant using the Nucleo-Mag Vet kit (Macherey Nagel). Viral loads in swab and tissue samples were determined using a previously published real-time RT-PCR protocol for measuring genomic viral RNA using a probe and oligonucleotide primers targeting the SARS-CoV-2 E gene.^50^

### Plaque titration assay

To quantify the infections virus titers, the inferior, middle and post-caval lobes of the lung and the distal half of the right brain hemisphere were homogenized in Tube M (Miltenyi Biotech, ref. 130-096-335) containing 1 ml of DMEM using a gentleMACS™ Tissue Dissociator (Miltenyi Biotech). Viral titers were determined by plaque assay on Vero E6 or Vero-TMPRSS2 cells as previously described.^37^ Briefly, 1×10^5^ cells/ml were seeded in 24-well plates 1 day prior to titration. Cells were inoculated with 100 μl of organ homogenate and the inoculum was serially diluted in cell culture medium at 1:10 dilution steps. The inoculum was removed 1 hour post-infection and replaced with overlay medium consisting of DMEM supplemented with 1.2% Avicel (RC-581, FMC biopolymer), 15 mM HEPES, 10% heat-inactivated FBS, 100 μg/ml streptomycin, and 100 IU/ml penicillin. Cells were incubated for 48 hours at 37°C and 5% CO_2_, fixed with 4% (v/v) neutral buffered formalin, and stained with crystal violet.

### Histopathological and immunohistochemical analysis

The left lung and the left hemisphere of the brain were collected upon necropsy either at day 5 or 14 pi with SARS-CoV-2-S^D614G^ and immersed in 10% neutral-buffered formalin. Following fixation, both tissues were embedded in paraffin, cut at 4 μm, and stained with haematoxylin and eosin (H&E) for histological evaluation. Lung tissue pathology was scored according to a previously published scoring scheme (**Supplementary Table 1**).^51^ A 1:3000 dilution of a rabbit polyclonal anti-SARS-CoV nucleocapsid antibody (Rockland, Cat. no. 200-401-A50) was used for SARS-CoV-2 immunohistochemical (IHC) analysis of the lung and brain. Paraffin blocks were cut at 3 μm, placed in a BOND RXm immunostainer (Leica Biosystems, Germany), and incubated for 30 minutes with the first antibody at room temperature. Antigen retrieval was performed by incubating the slides with citrate buffer for 30 min at 100°C. Subsequently, BondTM Polymer Refine Detection visualisation kit (Leica Biosystems, Germany) was used for signal detection using DAB as chromogen. The tissue sections were counterstained with hematoxylin.

### Statistical Analysis

Statistical analysis was performed using the GraphPad Prism program package version 8. Depending on the number of data groups to be compared, significant differences were calculated with either the Student’s t test (2 groups), the one-way ANOVA test (>2 groups, one time measurement), or the two-way ANOVA test (>2 groups, different time points). Dunnet’s multiple comparison was used for the one-way ANOVA test if each group was compared with a control group. Tukey’s multiple comparison was used for the two-way ANOVA test if different groups were compared at the same time point). Sidak’s multiple comparison was used for the two-way ANOVA test if different time points within one group were compared. Y values were transformed to Y=log(Y) or Y= log2(Y) if log_10_ or log_2_ logarithmic scales were used. If not otherwise stated, all tests were performed two-sided. The one-sided Mann-Whitney test was used to compare tetramer staining of CD8+ cells from naïve mice with the tetramer staining of CD8+ cells of either of the vaccine groups.

### Data availibility

All data generated or analysed during this study are included in this published article (and its supplementary information files).

## Acknowledgements

We thank Katarzyna Sliz, Daniel Brechbühl, Isabelle Wymann, Jan Salchli, Roman Troxler and Hans-Peter Lüthi for dedicated animal care. This work was supported in part by the Swiss National Science Foundation grants 31CA30_196062 (CB), 310030_189151 (GZ), and NRP78 4078P0_198473 (VT), COVID-19 special funds from the Swiss Federal Office of Public Health and the Swiss Federal Food Safety and Veterinary Office. We thank Artur Summerfield for help in securing Swiss government funds for this project. We thank the NIH Tetramer Core Facility for providing SARS-CoV-2 S-tetramers. We are grateful to Markus Hofmann for providing the Vero-TMPRSS2 cells.

## Author contributions

Conceptualization and project design: GZ, CB; investigation and data acquisition: AT, GZ, IBV, CD, RB, CB, GK, SW, PP; supervision, data analysis and interpretation: GZ, AT, CB; funding acquisition: CB, GZ, VT. All authors read and approved the paper.

---

**Supplementary Table 1.** Histopathologic assessment of lung pathology.

| Evaluation criteria | Score<sup>1</sup> |  |  |  |  |
| --- | --- | --- | --- | --- | --- |
|  | 0 | 1 | 2 | 3 | 4 |
| Distribution of lesions | Absent | Focal | Multifocal<br>(to coalescing) | Diffuse |  |
| Affected tissue (%) | <1% | 2-10% | 11-25% | 26-50% | >50% |
| Expansion to peripheral lung regions | No | Yes |  |  |  |
| Pulmonary atelectasis | Absent | Minor | Major |  |  |
| Peribronchial inflammation | Absent | Minor | Major |  |  |
| Perivascular inflammation | Absent | Minor | Major |  |  |
| Interstitial inflammation | Absent | Minor | Major |  |  |
| Intra-alveolar inflammation | Absent | Minor | Major |  |  |
| Bronchial epithelial necrosis | Absent | Minor | Major |  |  |
| Alveolar wall necrosis | Absent | Minor | Major |  |  |
| Infiltration by neutrophils | Absent | Minor | Major |  |  |
| Infiltration by macrophages | Absent | Minor | Major |  |  |
| Infiltration by lymphocytes | Absent | Minor | Major |  |  |
| Alveolar edema | Absent | Minor | Major |  |  |
| Perivascular edema | Absent | Minor | Major |  |  |
| Fibrinoid degeneration of vascular walls | Absent | Minor | Major |  |  |
| Vascular thrombosis | Absent | Minor | Major |  |  |
| Vasculitis | Absent | Minor | Major |  |  |
| Hemorrhage (interstitial, intraalveolar) | Absent | Minor | Major |  |  |
| Pleuritis | Absent | Minor | Major |  |  |

<sup>1</sup> The scoring of lung lesions was performed according to Dietert et al, 2017.<sup>51</sup>

**Supplementary Fig. 1.**
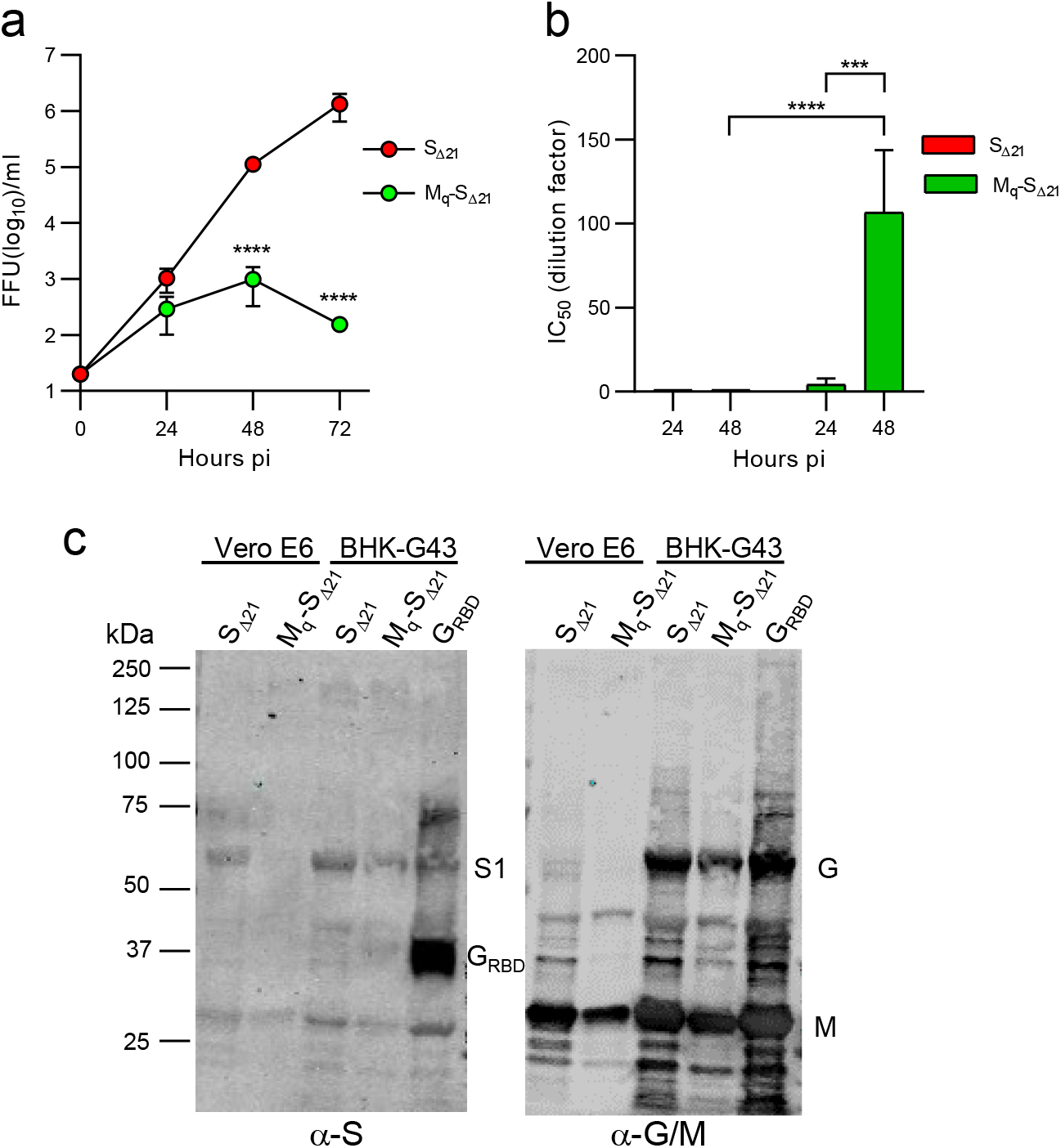
Attenuation of VSV*M_q_ΔG-S_Δ21_ by the induction of type I interferon. A549-hACE2/hTMPRSS2 cells were infected with either VSV*ΔG-S_Δ21_ or VSV*M_q_ΔG-S_Δ21_ using an moi of 0.01 ffu/cell. (**a**) Infectious virus produced at the indicated time points was determined by titration of the cell culture supernatants on Vero E6 cells. (**b**) Quantification of antiviral activity in conditioned medium on the VSV*ΔG(FLuc) reporter virus. Mean values and SD were calculated from 3 infection experiments. Statistically different values were computed using the two-way ANOVA test (***p = 0.0005; ****p < 0.0001). (**c**) Analysis of recombinant VSV-vectored vaccines by Western blot after SDS-PAGE under reducing conditions. See also the legend to Fig. 1 for details.

**Supplemental Fig. 2.**
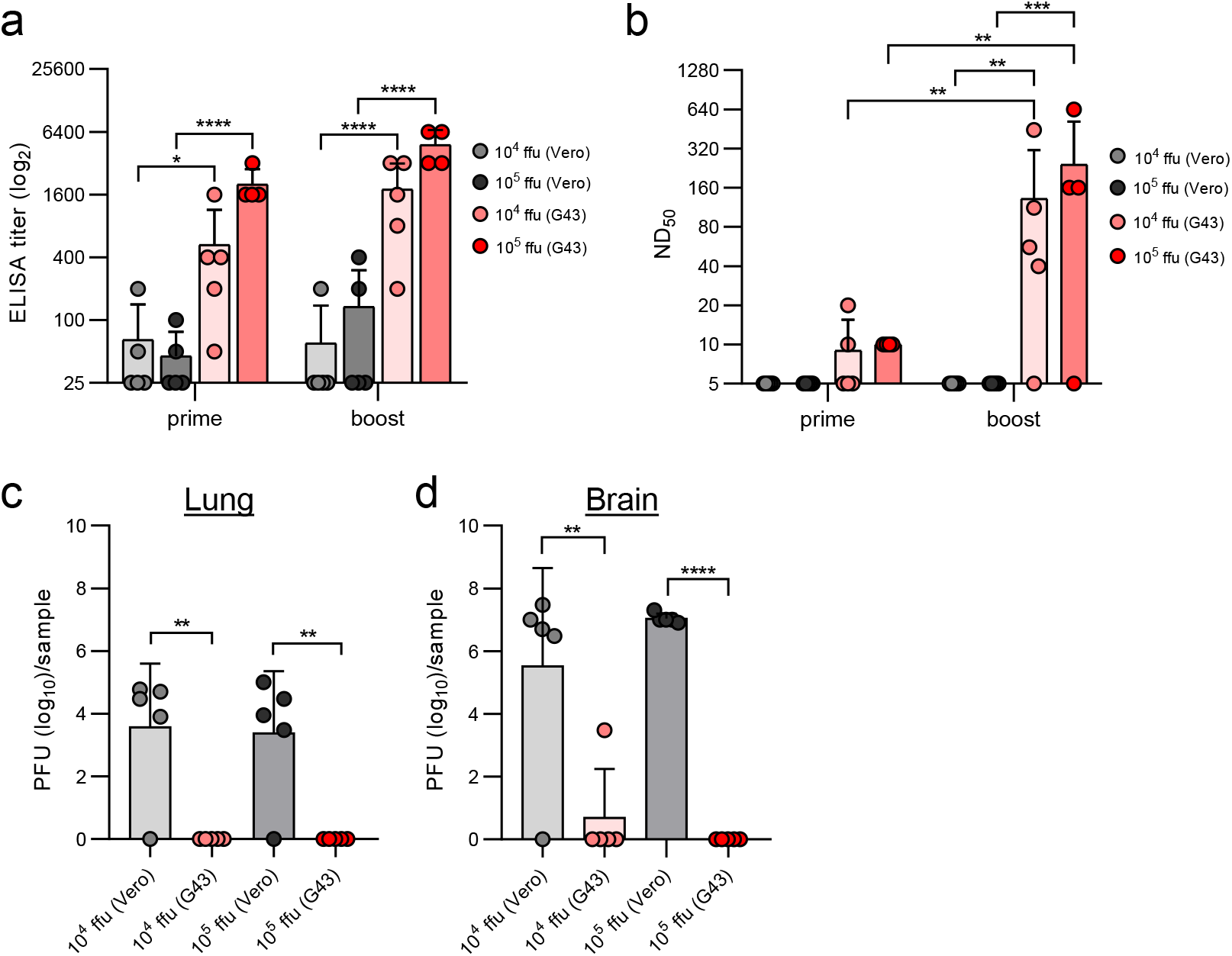
Comparison of ACE2-dependent and -independent immunization of C57BL/6 mice. C57BL/6 mice (group size n = 5) were immunized (i.m.) with either 10^4^ or 10^5^ focus-forming units (ffu) of VSV*ΔG-S_Δ21_ vector produced on either Vero E6 or BHK-G43 cells. (**a**) Detection of spike-specific serum antibodies by ELISA 3 weeks after the first (prime) and 3 weeks after the second immunization (boost). (**b**) Determination of the virus neutralization dose 50% (ND_50_) in serum of immunized mice. (**c, d**) Determination of infectious SARS-CoV-2-S^D614G^ titers in homogenates of lung (**c**) and brain homogenates (**d**) at 9 days pi. Mean values and SD are shown in all data sets. Significantly different values were calculated by the two-way ANOVA test (**b**) and by the one-way ANOVA test (**c, d**) (*p < 0.05, **p < 0.01; ***p < 0.0005, ****p < 0.0001).

**Supplemental Fig. 3.**
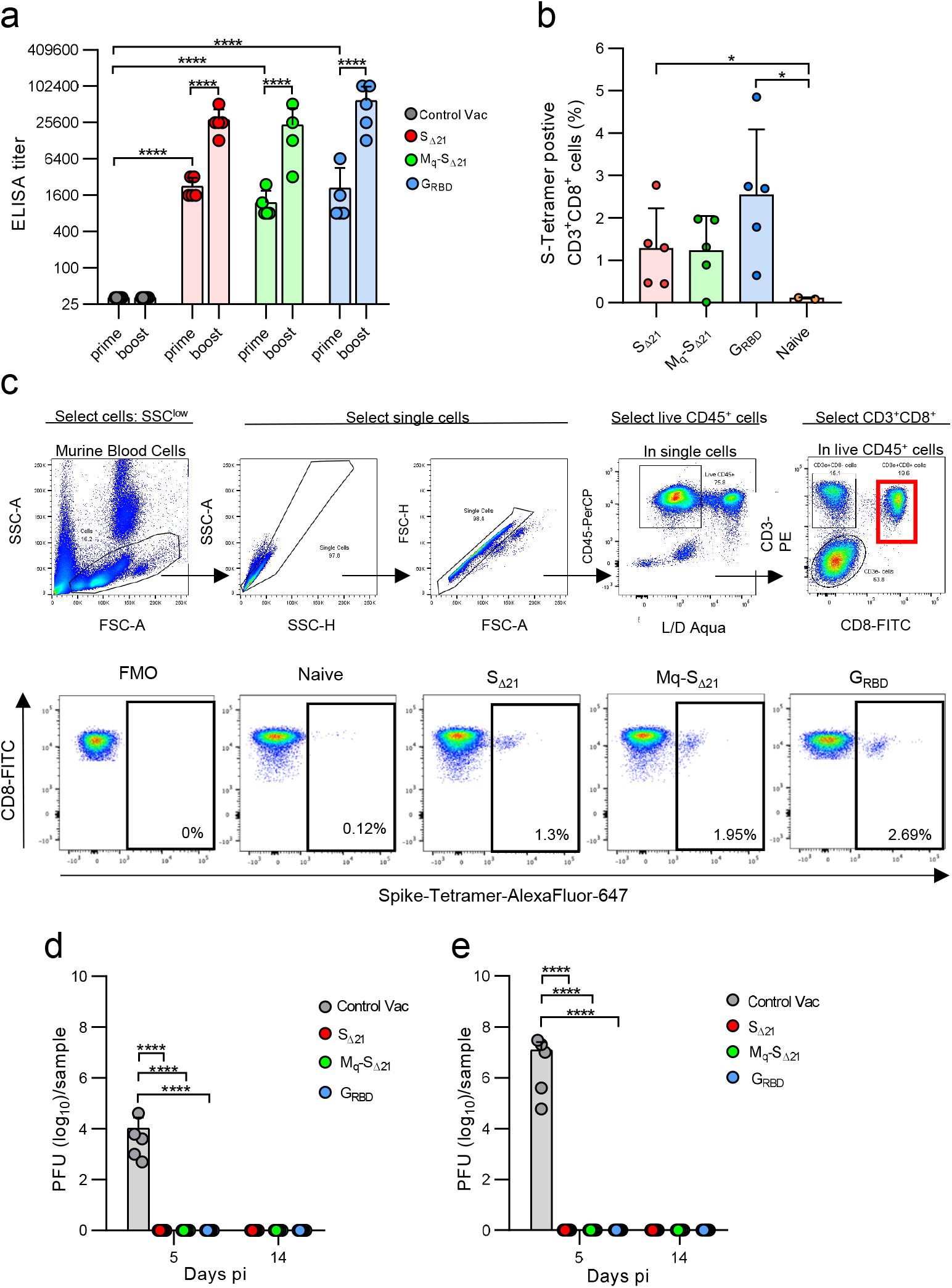
Humoral and cellular immune responses of vaccinated K18-hACE2 mice. (**a**) Detection of spike-specific serum antibodies by ELISA 3 weeks after the first (prime) and 3 weeks after the second immunization (boost) in the indicated vaccine groups (n = 5). (**b**) Frequency of spike protein-specific cells in the CD3^+^/CD8^+^ blood lymphocyte population 3 weeks after the second immunization with the indicated vaccines. (**c**) Representative flow cytometric analysis of CD3^+^/CD8^+^ lymphocytes after staining with the spike-specific tetramers. Upper panel: Gating strategy. Lower panel: Representative tetramer staining of lymphocytes from a single animal per each vaccine group (lower panel). (**d, e**) Determination of infectious SARS-CoV-2 titers by plaque assay in lung (**d**) and brain (**e**) tissue homogenates prepared from animals euthanized at either day 5 or day 14 pi. Mean values (represented by bar heights) and SD are shown. Statistically significant values were determined by the two-way ANOVA test (**a**), the one-tailed Mann-Whitney test (**b**), and the one-way ANOVA test (**d, e**) (*p < 0.05, ****p < 0.0001).

**Supplementary Fig. 4.**
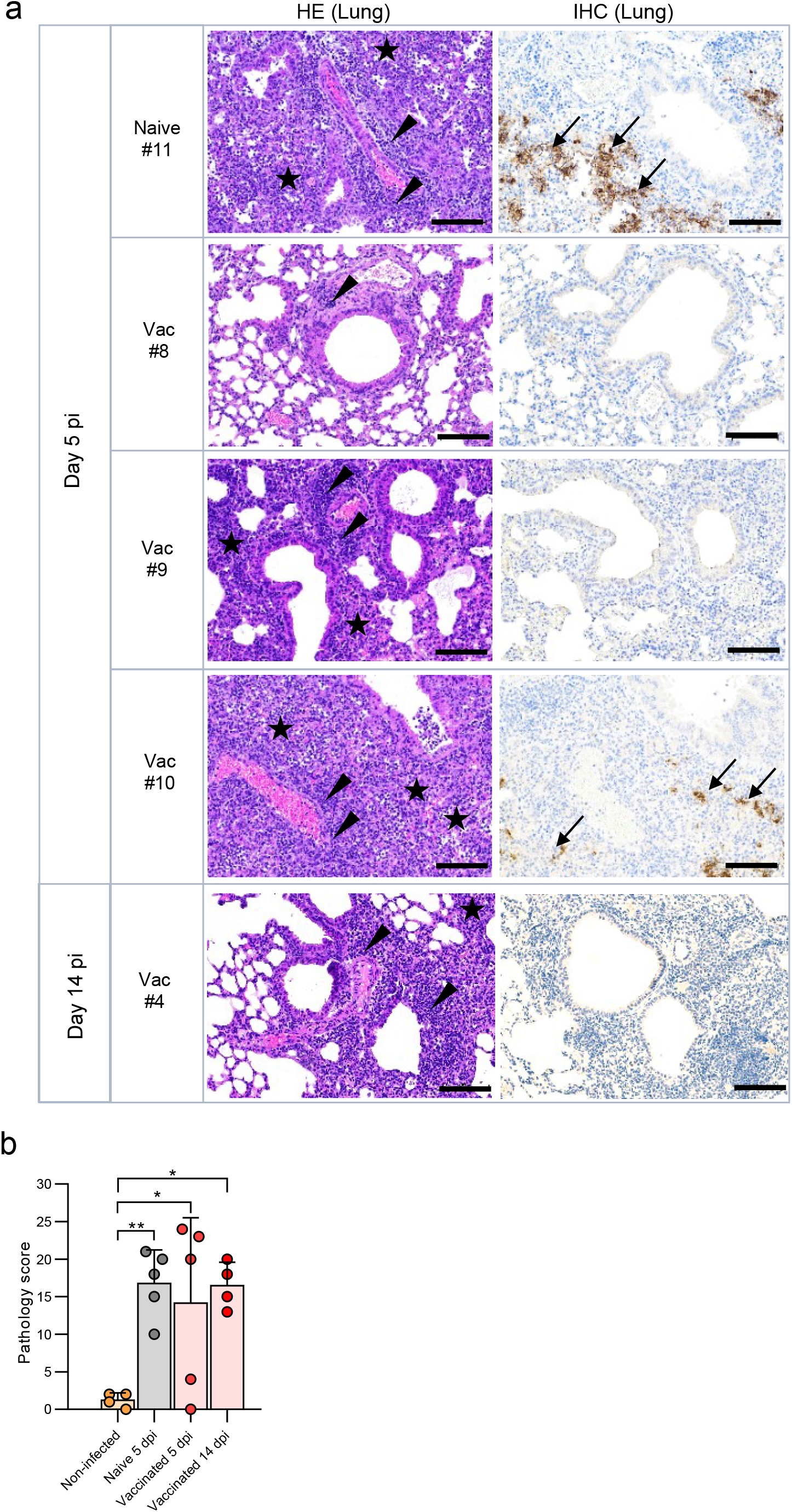
Histological analysis of lung tissue sections following SARS-CoV-2 Delta VOC infection. (**a**) Naïve and vaccinated K18-hACE2 mice were euthanized at day 5 and day 14 pi with SARS-CoV-2^Delta^ VOC and lung sections from individual mice analyzed by HE staining (HE Lung) and viral nucleoprotein immunostaining (IHC Lung). Arrow heads indicate perivascular/peribronchiolar infiltration. Stars mark tissue consolidation. Arrows indicate areas where SARS-CoV-2 nucleoprotein antigen was detected by IHC. The animal ID numbers shown on the left side of the figure identify the animals labeled in Fig. 7c and Fig. 7g. Bars = 100 µm. (**b**) Histopathological scoring of lung tissue sections prepared from vaccinated and naïve mice 5 and 14 days pi. Mean values and SD are indicated. Statistical differences were calculated with the one-way ANOVA test (*p < 0.05; **p < 0.01).

**Supplementary Figure 5:**
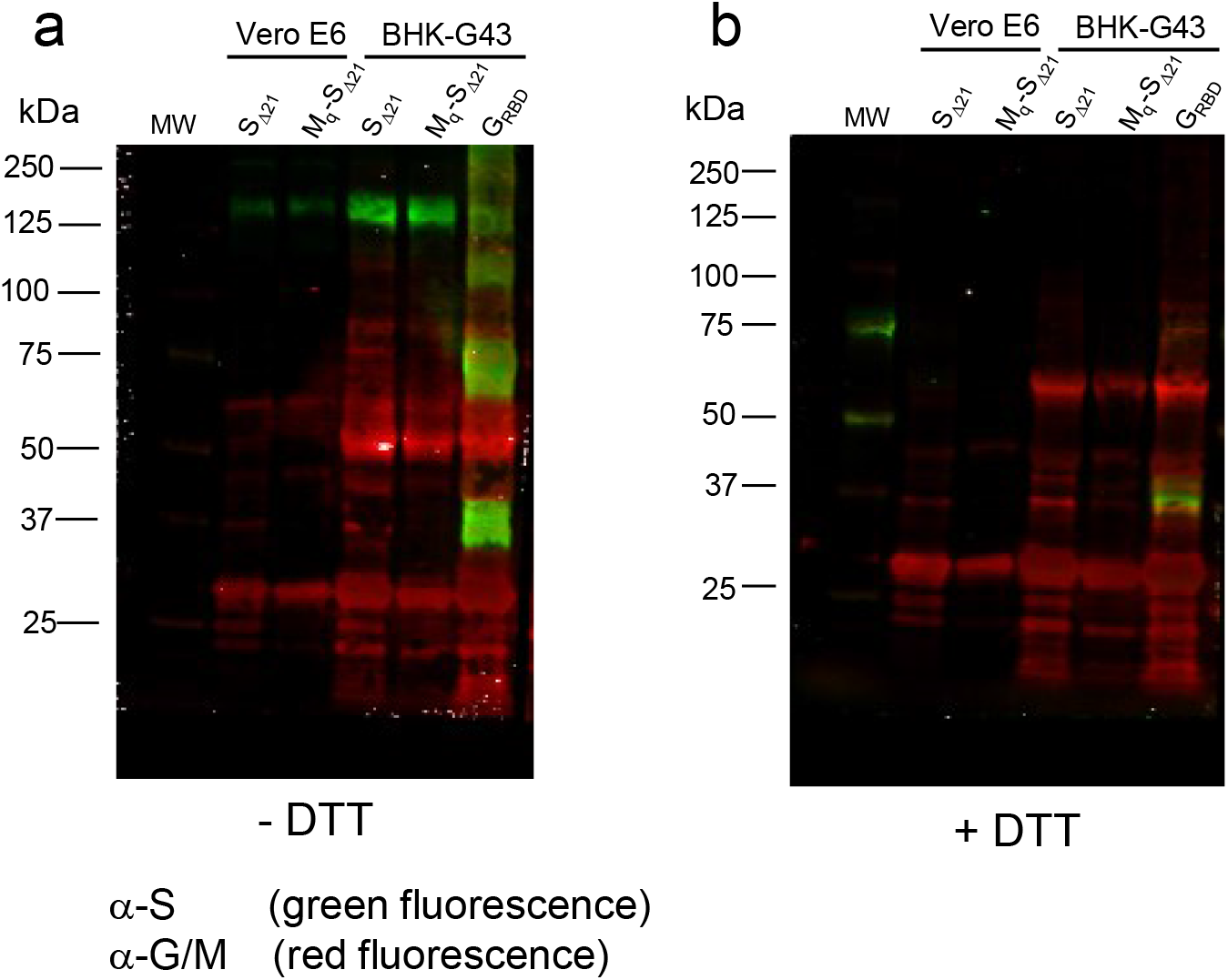
Western blot analysis of recombinant VSV vector particles. VSV*ΔG-S_Δ21_ (S_Δ21_) and VSV*M_q_ΔG-S_Δ21_ (M_q_-S_Δ21_) were propagated on both Vero E6 and BHK-G43 cells, while VSV*ΔG-G_RBD_ (G_RBD_) were propagated only on BHK-G43 helper cells. At 24 hours pi, the virus particles were concentrated from the cell culture supernatant by ultracentrifugation and dissolved in SDS sample buffer. The viral proteins were separated by SDS-PAGE under (**a**) non-reducing (-DTT) or (**b**) reducing (+ DTT) conditions and blotted onto nitrocellulose membrane. Antigens were detected with a COVID-19 convalescent serum (α-S) and a rabbit polyclonal immune serum directed to the VSV G and M proteins (α-G/M). The human convalescent serum was detected with a goat anti-human IRDye 800CW serum (green fluorescence), while the rabbit immune serum was detected with a goat anti-rabbit IRDeye 680RD serum (red fluorescence). The blots were derived from the same experiment and were processed in parallel.

